# Pangenome discovery and characterization of human protein-coding duplicated genes

**DOI:** 10.64898/2026.08.05.743125

**Authors:** Luyao Ren, DongAhn Yoo, Katarina Vlajic, Philip C. Dishuck, Xavi Guitart, Youngjun Kwon, Jiadong Lin, Katherine M. Munson, Kendra Hoekzema, Human Genome Structural Variation Consortium (HGSVC), Human Pangenome Reference Consortium (HPRC), Andrew B. Stergachis, Mitchell R. Vollger, Devin K. Schweppe, Evan E. Eichler

## Abstract

Protein-coding genes mapping to high-identity segmental duplications (SDs) have been difficult to annotate and characterize and are the source of most previously unknown protein-coding genes being discovered as part of the human pangenome. Here, we combine long-read assembled human genomes (298) and long-read transcriptome data (5.6 billion full-length cDNA from 83 tissues) to phylogenetically interrogate 493 gene families discovering 2713 potentially copy number polymorphic genes not present in the human reference genome. For reference SD gene families where paralog specificity can be assigned, we find that 60.0% are expressed and maintain open reading frames, with 45.7% showing high expression in brain, embryo, or testis. We revise 386 gene models, including 150 that absent or different from current T2T-CHM13 gene annotation and 236 (35.1%) pseudogenes as protein-coding where we find evidence of transcription, an open reading frame, and chromatin-accessible promoters. We find that 24.2% of SD genes show evidence of constraint for both copy number and amino acid mutation. The majority of these constraint genes are ancestral, whereas only 16.2% of derived duplicated genes that emerged recently in the human lineage show evidence of constraint. The pangenome provides unparalleled specificity to understand genetic variation in SD genes allowing us to distinguish functional genes from pseudogenes and highlighting potential gene innovations that arose most recently in human evolution.

## INTRODUCTION

Segmental duplications (SDs) are large genomic regions (>1 kbp) that share high sequence identity (>90%)^1^. In humans, SDs are enriched for genes and show higher degrees of transcriptional diversity^2^. More than 1000 protein-coding genes annotated in the T2T-CHM13 reference overlap by >30% with sequence annotated as segmentally duplicated^3,4^ (Fig. 1). Because of their high sequence identity and their proclivity for non-allelic homologous recombination (NAHR) and interlocus gene conversion (IGC), SD genes are frequently copy number variable^5–7^ and show a 10-fold increase in structural variation when compared to unique regions^8,9^. It has been estimated, for example, that approximately half of all structural variants (SVs) larger than 1 kbp map to SDs, even though SDs comprise only ∼6% of the human reference genome^4,10^. This hypermutability leads to rapid evolutionary turnover of duplicated genes, characterized by a high birth-death rate with frequent pseudogenization, while also creating opportunities for the emergence of new genes with new functions^11^ and generating extensive copy number variation among different human haplotypes.

**Fig. 1.**
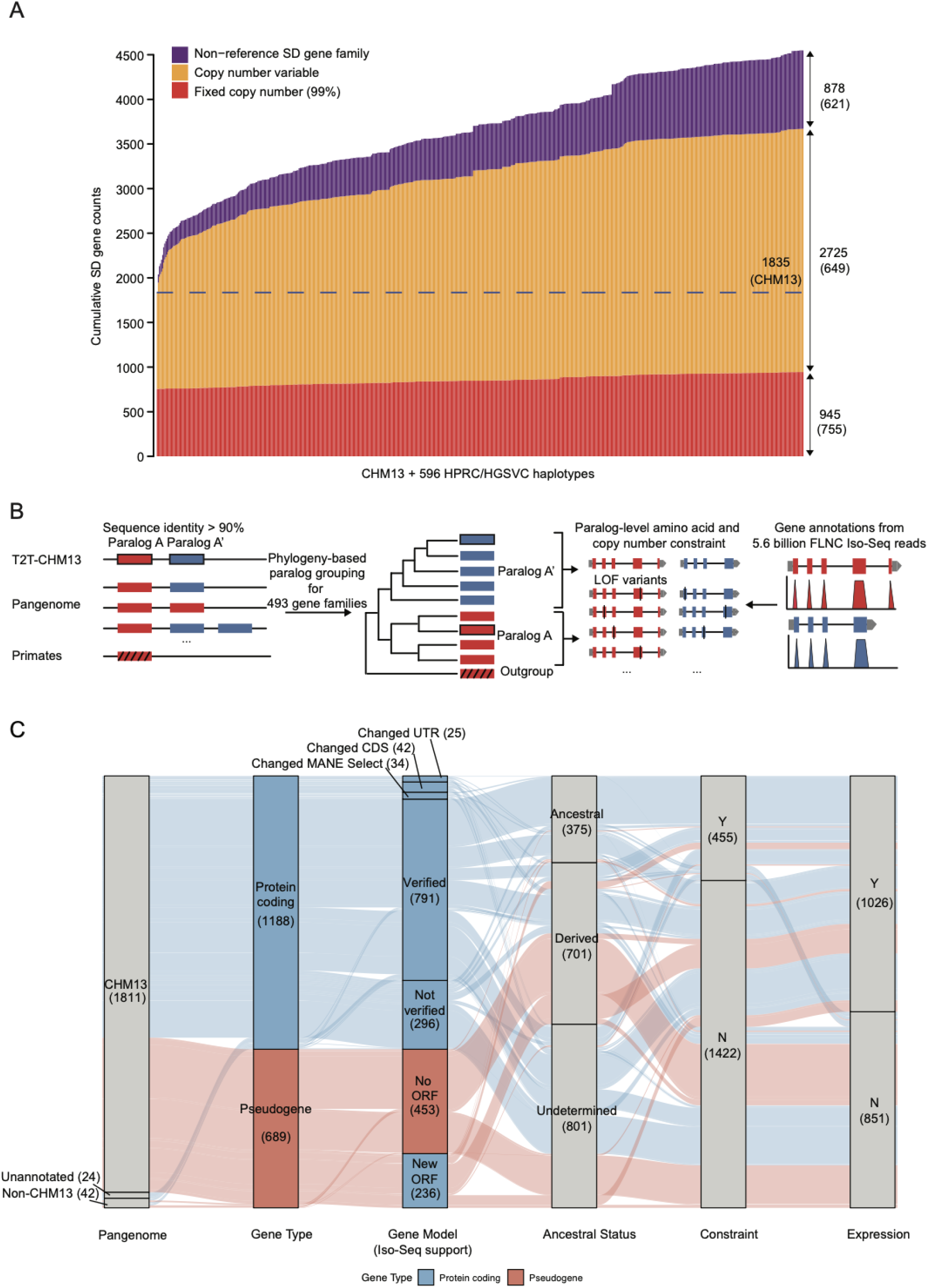
Pangenome and phylogenetic analysis of SD genes. **(A)** Cumulative curve of SD gene counts across the pangenome. The x-axis depicts the increase of SD genes by consecutive addition of 596 haplotype-resolved assemblies to T2T-CHM13 annotation. The y-axis shows the cumulative sum of maximum gene counts per SD gene across the assemblies. Red indicates fixed SD genes, defined as genes present in more than 99% of haplotypes at a stable copy number of one. Orange indicates copy number polymorphic SD genes. Purple indicates non-CHM13 SD gene families that are represented by a single, nonduplicated locus in T2T-CHM13 but are duplicated elsewhere in the pangenome, identified from a subset of 458 assemblies (Methods). The numbers on the right indicate the maximum copy number for each SD gene category, with the number of monophyletic clades shown in parentheses. There are a total 2713 duplicated genic loci added by the pangenome. **(B)** Study workflow overview. Intron sequence corresponding to existing gene models was extracted to build a multiple sequence alignment (MSA) and phylogeny (not location) was used to define paralogs which were assessed for amino acid and copy number constraint. Long-read transcriptome data were then used to annotate gene models and tissue expression. **(C)** The alluvial plot shows three gene categories: 1811 SD genes from 493 gene families annotated in the T2T-CHM13 reference, 24 protein-coding genes with sequence present in T2T-CHM13 but absent from its current annotation, and 42 non-CHM13 genes identified from phylogenetic trees as belonging to clades containing no T2T-CHM13 genes. Their refinement was based on RefSeq gene annotation, long-read Iso-Seq validation of gene models, ancestral status, evidence of copy number and amino acid constraint across the human pangenome, and long-read Iso-Seq expression. Gene models are assessed and verified by using a resource of 5.6 billion Iso-Seq reads from multiple data sources (Supplementary Table 4), with a minimum of five supporting reads, and then compared against T2T-CHM13 RefSeq annotation. The expression category used for tissue-specific expression analysis was defined more stringently, as transcripts per million (TPM) > 5 in at least one library or TPM > 1 in at least three libraries.

In humans, some of the most recent high-identity duplications have contributed to unique neuroadaptive features that have, in part, promoted expansion of the frontal cortex. For example, more than half a dozen neofunctional genes have emerged as a result of human-specific SDs. Functional studies have shown that these genes play roles in increasing synaptic connectivity (*SRGAP2C*^12^), increasing the number of basal radial progenitor cells (*TBC1D3*^13^ and *ARHGAP11B*^14^), modifying cortical neurogenesis through mTOR signalling (*CROCCP2*^15^), and promoting protracted neural development (*NOTCH2NL*^16^). Despite their importance, however, these neofunctional genes were either missing or misannotated in the early drafts of the human genome because the high-identity paralogs were misassembled or because gene-annotation tools imposed the architecture of the ancestral genes onto the derived duplicate genes whose gene structure had frequently changed as part of their recent evolution^16–21^. Moreover, the characterization of duplicated genes has lagged far behind unique genes because of the challenges of reliably assigning short-read sequence data to these ∼1000 copy number polymorphic, protein-coding genes in high-identity SDs^22^. Thus, they have been systematically excluded from population-genetic databases, expression studies, and disease-association studies^23,24^, despite their critical role in disease and evolution.

As an application of the human pangenome resource^25^, we address this longstanding limitation of the human genome and systematic gap in characterizing these genes by leveraging both long-read genome and transcriptome data. We use the genetic diversity and complete, highly accurate SD sequences represented in a total of 298 genomes from the Human Pangenome Reference Consortium (HPRC)^25^ and the Human Genome Structural Variation Consortium (HGSVC)^26^ to phylogenetically classify and interrogate variation of each human duplicated gene family at the level of the individual paralog. We then integrate long-read transcriptome data to not only assess the expression pattern of each duplicate but also to reannotate the predominant human protein-coding gene models for each member of the gene family (Fig. 1B-C). These data are used to develop an initial mutational constraint map for both copy number and amino acid changes at the paralog-specific level, helping distinguish candidate functional copies from pseudogenes. Our analysis reveals hundreds of new gene models providing a roadmap for pinpointing the most functional duplicate protein-coding genes. We also highlight potential gene innovations where the gene structure has changed and reclassify some longstanding pseudogenes as expressed copies that maintain an open reading frame (ORF) where both epigenetic data and limited proteomic data suggest the presence of *bona fide* transcription and translation into proteins.

## RESULTS

### Pangenome analysis and phylogeny-based grouping

Based on existing RefSeq annotations for completed genome (T2T-CHM13; [JHU RefSeqv110 + Liftoff v5.2](https://github.com/marbl/CHM13)), we identify 1811 “genic loci” mapping to SDs and assign these to 493 SD gene families (Fig. 1C, Supplementary Table 1). The set includes 1139 protein-coding genes and 672 pseudogenes, including genes predicted to no longer be transcribed or have coding potential (Methods). Using T2T-CHM13 as a baseline, we analyzed SD content for 596 assemblies (298 samples x 2 haplotypes) generated from HPRC and HGSVC. We identify 945 SD genes as fixed (present in >99% of haplotypes at a stable copy number of one) and 2725 as copy number polymorphic. In a subset of 458 assemblies with available de novo CAT gene annotations^25^, we validated SD regions using short-read sequencing read depth and identified 12% of the human genome (382.7 Mbp) as segmentally duplicated. From this subset, we further identify 878 non-reference SD gene families that are unique (not duplicated) in T2T-CHM13 but duplicated elsewhere in the pangenome (Fig. 1A, Extended Data Fig. 1, Table 1, Glossary).

**Table 1.**
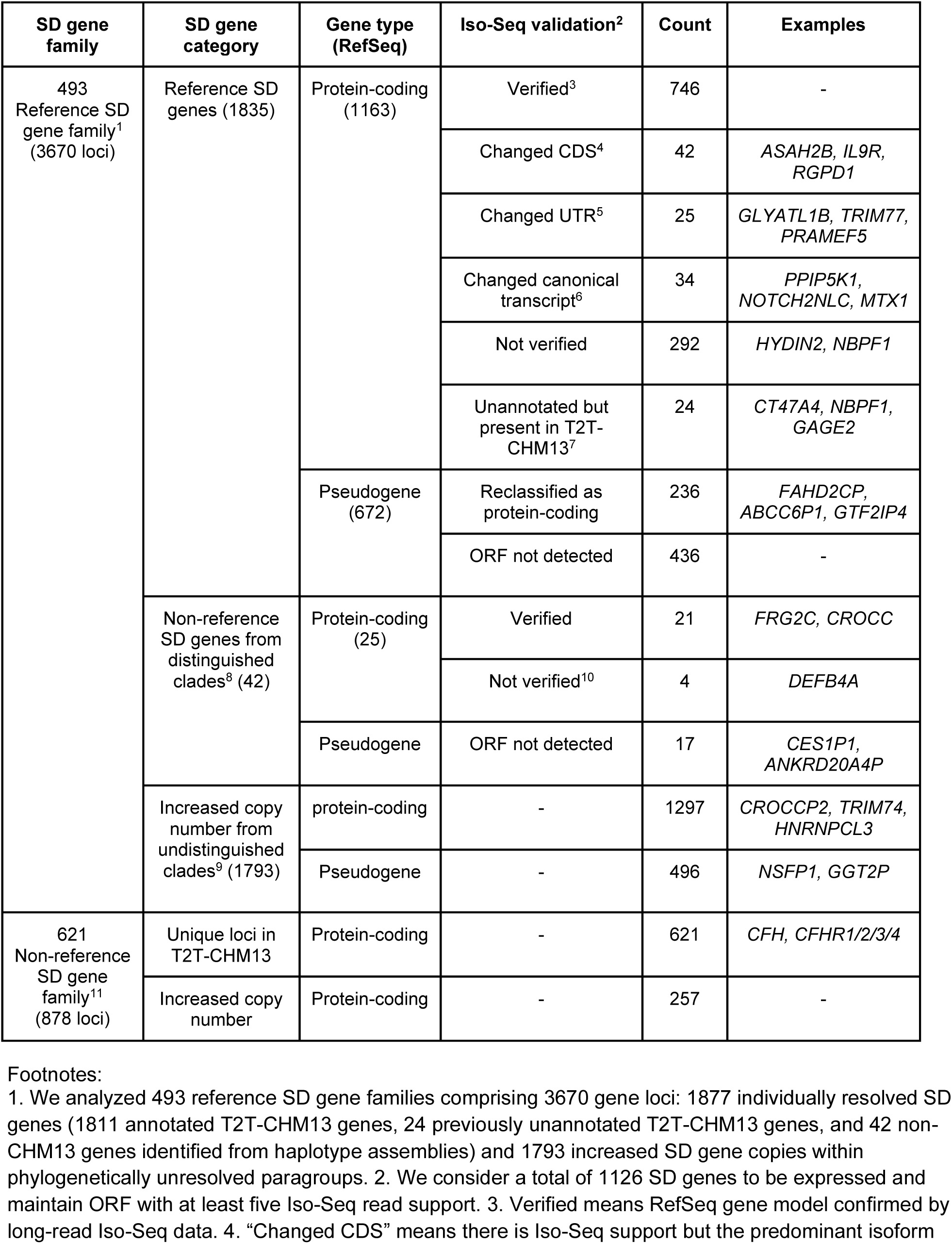
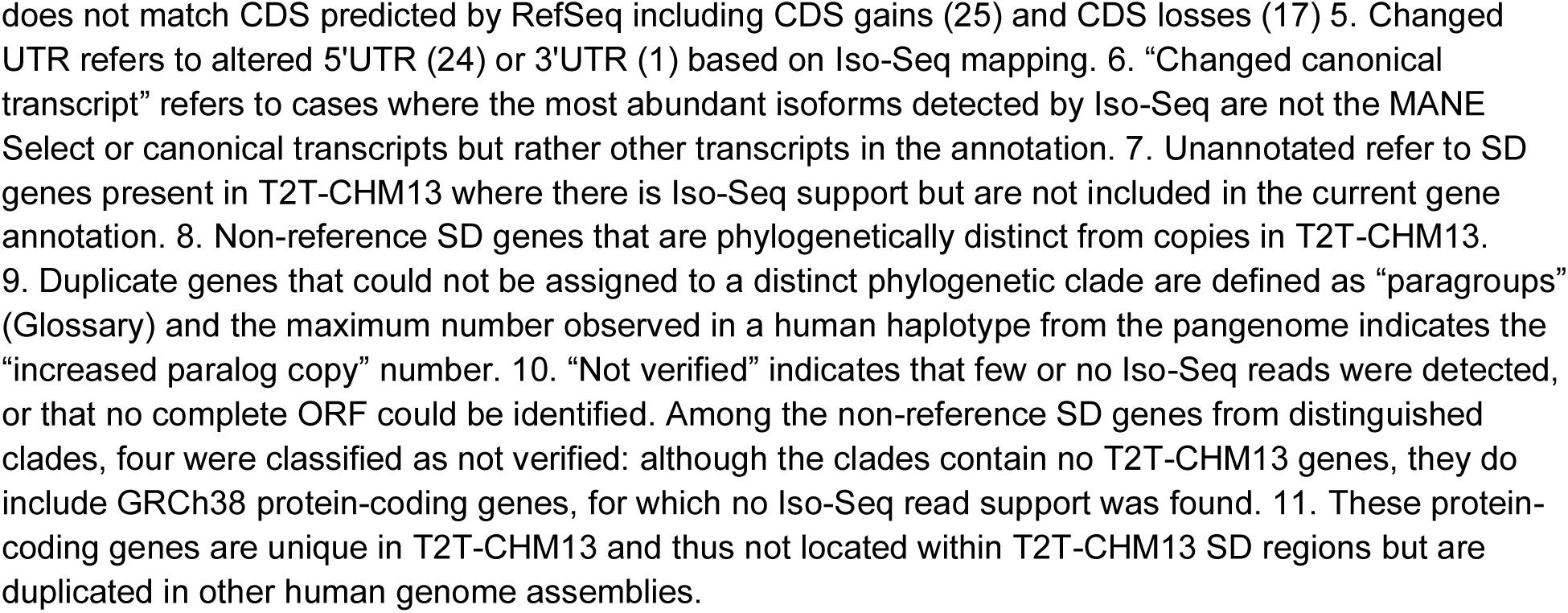
Summary of Iso-Seq validation of protein-coding duplicated gene models.

Focusing on the 493 protein-coding SD gene families in the T2T-CHM13, we apply a recent phylogenetic-based pangenome approach^20^ to group and distinguish highly identical paralogs for each of the haplotype-resolved genomes from the complete set of all 298 HGSVC and HPRC samples (Fig. 1B, Supplementary Fig. 1, Methods). For each gene family, we identify the corresponding genic loci from 596 haploid human genome assemblies that passed QC and construct multiple sequence alignments based on neutrally evolving intronic sequences from all allelic and paralogous copies. We then generate population-based phylogenetic trees for each of the 493 SD gene families, using recently completed nonhuman primate loci as outgroups^27,28^.

The tree topology serves three purposes. First, it organizes human genetic diversity of SD genes based on the natural history/evolution of each gene family providing an estimate when each paralog emerged over human/primate evolution based on the coalescence of human haplotypes (Fig. 2A). Second, it provides a framework for assessing both copy number changes as well as synonymous and amino acid changes occurring among different haplotypes or groups of closely related genes (Fig. 2B-C). Third, tree substructure provides a powerful tool for pinpointing potential sites of gene conversion as such events are likely to either “relocate” a paralog to a new syntenic position^21^ or result in hybrid/gene fusion with respect to the reference frequently characterized by an internal topology substructure (e.g., *CYP2D6* vs. *CYP2D7* [Supplementary Fig. 2] and *SMN1* vs. *SMN2* [Supplementary Fig. 3]).

**Fig. 2.**
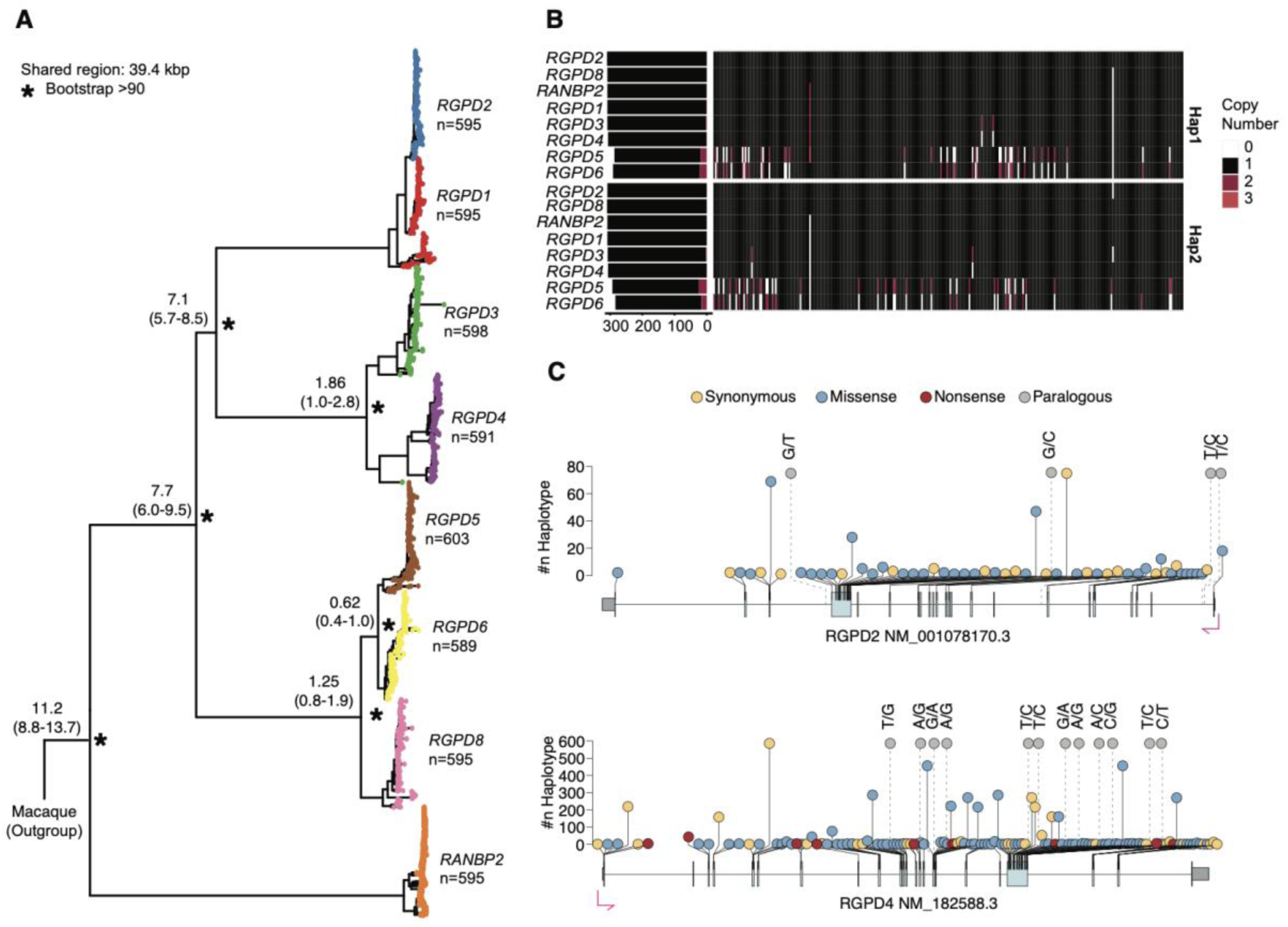
Paralog grouping and variant detection within the *RGPD* gene family. **(A)** Population-based phylogenetic tree constructed from HPRC and HGSVC phased genome assemblies for the *RGPD* gene family, with macaque (MFA) used as the outgroup. The tree was constructed from an MSA of 39.4 kbp of intronic (neutral) sequence. Colored dots distinguish different paralogs within the *RGPD* gene family annotated based on T2T-CHM13 and branches (*) indicate >90% bootstrap support. Node labels indicate the median divergence time (million years ago) with 95% confidence intervals (CIs). Integer values represent copy numbers in 596 human haplotypes and monophyletic substructure and values lower and greater than 596 denote interlocus gene conversion or copy number variation. **(B)** Heatmap summarizing the copy number of each paralog across both haplotypes for each reevaluated and reassigned individual HPRC/HGSVC haplotype, with gene deletions (white) and duplications (red) indicated. **(C)** Amino acid constraint showing synonymous (yellow), amino acid substitutions (blue), and stop codon polymorphisms (red) predicted using Variant Effect Predictor (VEP)^32^ based on a gene model with long-read transcriptome support. Dashed lines indicate paralog-specific variants. We showed 12 of the 41 *RGPD4* paralogous variants located in the coding sequence (CDS) and all 4 *RGPD2* paralogous variants distributed across the whole gene body. This analysis was repeated for all remaining 492 gene families.

As an example, we highlight the *RGPD* (*RANBP2 Like And GRIP Domain Containing 1*) gene family—a recently expanded human gene family under positive selection^29^ and thought to be involved in nuclear pore activity^30^. It originated through the duplication and juxtaposition of two ancestral genes, *RANBP2* and *GCC2*^31^, and we estimate that it began to duplicate before and after African-great ape speciation with a subset of copies emerging 1-2 million years ago specifically in the human lineage (Fig. 2A). In general, each paralog forms a distinct clade with a subset showing strong bootstrap support (>90%). Exceptions include *RGPD1* and *RGPD2* where we find evidence of extensive IGC resulting in the formation of “hybrid genes” at high frequency (422/595) in the population (Supplementary Fig. 4). Notably, only *RGPD1*, *RGPD2*, and *RGPD8* exist in a one-to-one relationship within each of the 595 haplotypes. Thus, we regard each as being nearly fixed as they were present in >99% of haplotypes (Fig. 2B), while other copies show evidence of copy number variation. For specific pairs of duplicate genes, *RGPD3–RGPD4* and *RGPD5–RGPD6,* longer-range synteny afforded by complete haplotype resolution of the pangenome linear assemblies suggests whole-gene IGC. Although these pairs consistently add up to two copies per sample, the individual paralogs themselves are not fixed at the gene level (Fig. 2B).

It should be noted that for some higher-copy gene families, such as those associated with core duplicons (*TBC1D3*, *NPIP*, *GOLGA*, *NBPF*, *SPDYE*, and *LRRC37A*)^2^, individual loci could not always be distinguished because paralogous sequence divergence approximates or is less than allelic variation (Supplementary Figs. 5-10, Methods). In such cases, we combine paralogs into larger multi-member monophyletic groups (termed *paragroups*, Supplementary Fig. 11; see Glossary and Supplementary Fig. 1B for schematic). For these, we estimate the overall copy number in the human population—usually an approximate integer value of 596. In total, we identify 1404 monophyletic clades with gene annotations among these 493 SD gene families (Supplementary Table 2).

Relative to the T2T-CHM13 reference, we identify 42 phylogenetically distinct duplicated genes not present in the reference genome—that is to say these paralogous genes can be distinguished but are not present in the T2T-CHM13 and, thus, represent novel divergent genes present only in a subset of human haplotypes. Similarly, we identify 1793 potential duplicated protein-coding genes that show higher copy number than T2T-CHM13 but which are so identical they cannot be phylogenetically distinguished (Extended Data Fig. 2). These pangenomic SD genes are assigned to 760 paragroups. Combining these “non-reference SD gene” copies (Glossary), we estimate 1835 copy number polymorphic duplicated genes from the human pangenome that are not present in the T2T-CHM13 completed reference human genome. Finally, there is a subset of loci (621) that are unique in T2T-CHM13 but correspond to genes but are duplicated in a subset of other human genomes (Fig. 1A). The maximum copy number of these is 878 (Supplementary Table 3) and we refer to this subset as non-reference SD gene families (Glossary).

### Transcript redefinition of duplicated gene models

Duplicate gene annotations have been particularly problematic and often in error due to the limitations of short-read transcriptome data^33–35^. Because ancestral copies are frequently highly conserved among vertebrates or primates, there is a preferential bias for ancestral unique gene or initial RefSeq gene models to be superimposed among other paralogs^21^. To assess the validity of gene annotations from RefSeq ([JHU RefSeqv110 + Liftoff v5.2](https://github.com/marbl/CHM13)), we compiled a database of full-length non-chimeric (FLNC) cDNA generated from 5.6 billion Iso-Seq reads from 583 libraries, representing 83 human tissue/cell types^21,36,37^ (Supplementary Table 4). We extracted Iso-Seq reads aligning to each paralog, reconstructed transcript models, and predicted their ORFs essentially leveraging the specificity of long-read sequencing (LRS) transcriptomic data to assign transcripts to individual paralogs or paragroups. For each paralog, we compare the most abundant and longest Iso-Seq isoforms to the canonical transcripts annotated in T2T-CHM13 RefSeq annotation (Methods).

We assessed all 1811 RefSeq gene/pseudogene models using Iso-Seq reads. Among the 1139 protein-coding genes in the reference annotation, we confirm 746 (65.5%) as valid by long-read transcriptome data (Table 1). We find 101 protein-coding gene models that show significant gene model discrepancies, including 42 CDS and 25 untranslated region (UTR) differences, and 34 cases in which RefSeq isoforms were supported, but the most abundant Iso-Seq isoforms differed from the canonical RefSeq annotation. For example, the Iso-Seq gene models for *RGPD1* (Extended Data Fig. 3B) and *ZNF587B* (Extended Data Fig. 3C) show an extended 5′ end, supported by reduced methylation and increased chromatin accessibility at the transcription start site (TSS)— i.e., the most likely location of the promoter. For *NOTCH2NLC*, the most abundant Iso-Seq transcript contains a fully untranslated first exon that is supported by 57.1% of reads (Supplementary Fig. 12A). Similarly, the predominant *PDPR_2* isoform, supported by 49.6% of reads, lacks two coding exons relative to the canonical RefSeq transcript (Supplementary Fig. 12B). For both genes, the longstanding canonical RefSeq transcript models represent fewer than 0.1% of long-read transcripts. We also identified 24 unannotated protein-coding genes in T2T-CHM13 that have Iso-Seq support but are absent from RefSeq annotations.

Using this Iso-Seq approach, we also reassessed 672 annotated pseudogene models and find that a remarkable 35.1% (236/672) show evidence of spliced transcript expression, retain an intact ORF, and are supported by at least five Iso-Seq reads (Supplementary Table 5). For example, *WASH5P_1*—a putative noncoding lncRNA whose overexpression is frequently associated with favorable cancer outcome^38^—not only maintains an ORF but also shows a novel protein-coding gene structure (Extended Data Fig. 3D).

To assess the likelihood that these new annotations represent *bona fide* protein-coding transcripts, we first evaluated whether they are predicted to undergo nonsense-mediated decay (NMD). Among these transcripts, 64 are predicted targets of NMD, based on CDS termination at least 50 bp upstream of the final exon–exon junction, leaving the majority 172 as non-NMD targets. Second, we searched for evidence of putative promoters using available HPRC lymphoblast long-read Fiber-seq datasets^25^. Specifically, we compared chromatin accessibility within ±500 bp of the TSS among three groups: transcriptionally inactive RefSeq pseudogene models (“inactive pseudogenes”) (n=436), pseudogene-derived loci (n=236) for which we defined new ORF-containing gene models (“resurrected genes”), and known protein-coding genes (n=1139) as a positive control (Methods). Resurrected genes show a significant enrichment of chromatin accessibility signals at promoters compared with inactive pseudogenes lacking expression evidence in the 5.6-billion-FLNC database (OR = 3.54, p = 2.98 × 10^-267; Fig. 3A,B).

**Fig. 3.**
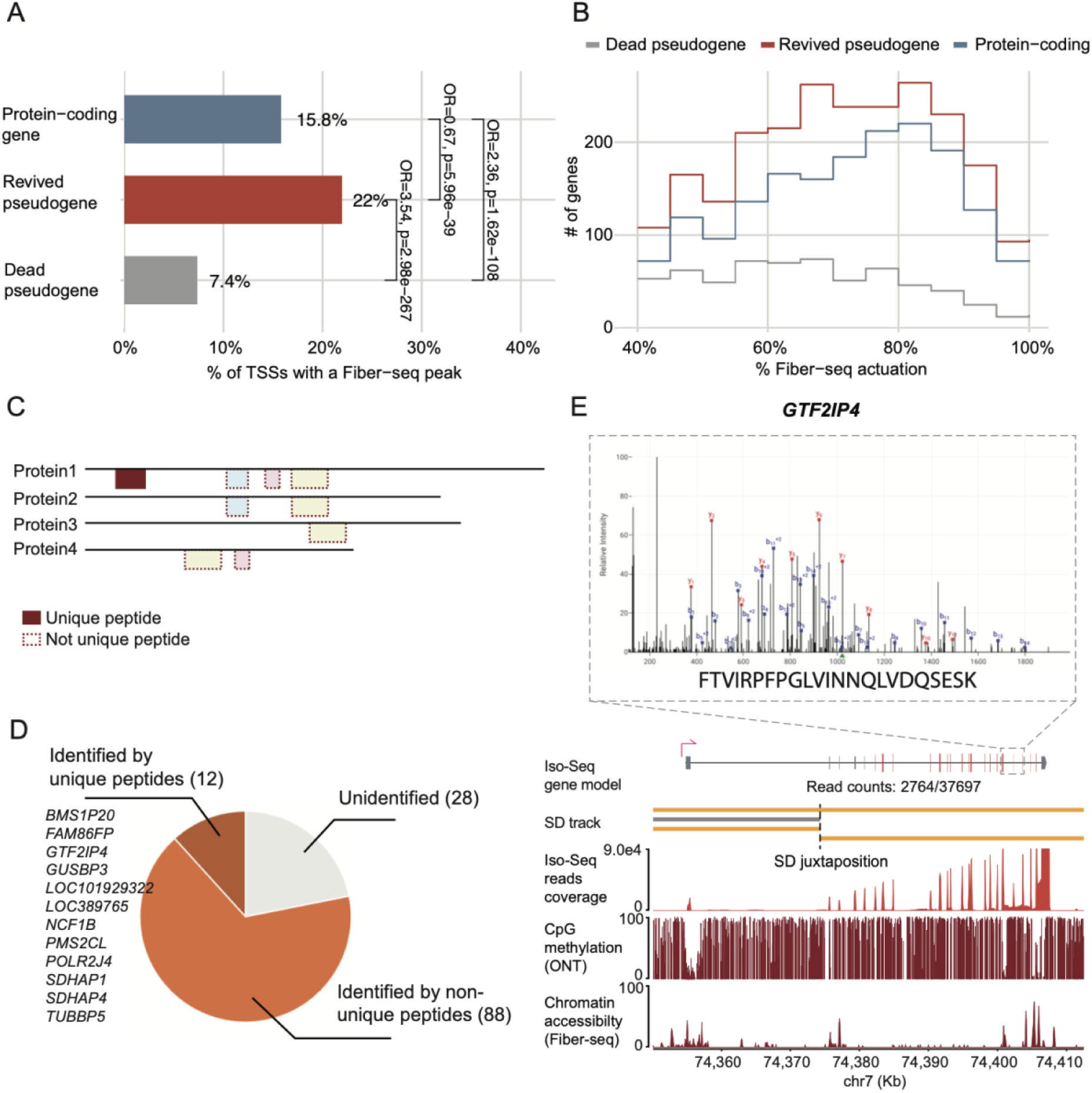
Fiber-seq and mass spectrometry support the reclassification of pseudogenes. **(A)** Accessibility at gene promoters (TSS +/-500 bp) in HPRC Fiber-seq data across three classes: dead pseudogenes (negative control), revived pseudogenes, and protein-coding genes (positive control). Fraction of loci in each class with at least one accessibility peak at the TSS. Brackets give Fisher’s exact test odds ratios and p-values for all three pairwise contrasts. Dead and protein-coding loci were subsampled to match the number of revived pseudogenes. **(B)** Distribution of maximum Fiber-seq chromatin actuation for loci with ≥40% accessibility. **(C)** Schematic plot showing unique and non-unique peptides. Non-unique peptides can be mapped to other known gene annotations in addition to the query gene, whereas unique peptides map only to the query gene annotation. **(D)** Number of reclassified pseudogenes validated by mass spectrometry. **(E)** Selected example of a reclassified pseudogene (*GTF2IP4*) supported by unique peptides. *GTF2IP4* was identified with eight peptide-spectrum matches (PSMs) across three unique peptides, detected exclusively in the medulloblastoma dataset. The three unique peptides supporting *GTF2IP4* identification comprised one high-confidence peptide (FTVIRPFPGLVINNQLVDQSESK, six PSMs at charge states 3+ and 4+) and two lower-confidence peptides derived from the same tryptic site: a fully cleaved 22-aa peptide and its missed-cleavage variant extending into the adjacent downstream tryptic fragment, each identified with a single PSM.

Finally, we searched for protein evidence from public human mass spectrometry-based proteomics data (Methods). Although peptide-level support can be detected for 78.1% (100/128) of these models where a peptide is predicted, we caution that due to high sequence similarity among duplicated genes, many detected peptides are shared between paralogs and, thus, cannot be assigned to a specific gene copy because the proteins are virtually identical (Fig. 3C). Only 12 pseudogene models are supported by unique peptides, providing the most stringent evidence for their protein-coding potential and locus-specific expression for these reclassified pseudogenes (Fig. 3D). Such is the case for *GTF2IP4* (Fig. 3E) —a member of the general transcription factor 2I mapping to the breakpoint region associated with Williams-Beuren syndrome and thought to be important for processes associated with myelination^39^. Combining the Iso-Seq, ORF, promoter, and NMD results, we conservatively estimate 172 additional protein-coding genes, adding 165.2 kbp coding sequence back to human genome annotation.

Because the reference genome T2T-CHM13 represents only one particular haplotype, we searched for evidence of new or additional duplicated loci with ORFs and therefore potential protein-encoding genes that would be polymorphic in the human population. Phylogenetic analysis identified 42 genetically distinct copies that could be distinguished from existing monophyletic clades. Of these, 25 are protein-coding, 14 are present in GRCh38, and 11 absent from both reference genomes are supported by Iso-Seq data. These loci represent recently duplicated candidate genes that have not yet reached fixation in the human population and can be directly evaluated for transcriptional support when they are sufficiently frequent in the population. Notable examples include a duplicated truncated version of CROCC (39.7 kbp) identified in 56.5% (337/596) haplotypes (Extended Data Fig. 3A) where we find evidence of strong transcript support as well as hypomethylation signal reduced evidence of a promoter.

SDs provide a genomic substrate for gene fusion because recurrent rearrangements can juxtapose SD blocks and bring otherwise separate genes into close proximity^40^. We identify 51 potential duplicated gene fusions produced by juxtaposed SD blocks of different origins, each with abundant read support (>5 reads) and that maintain at least one ORF (Supplementary Table 5). Of these, 16 retain predicted CDS from both individual genes. Seventeen of these fusions involve juxtaposed SDs that placed core duplicon genes downstream of partner-gene promoters and 5′ exons, generating fusion transcripts predicted to encode proteins in which the core duplicon defines either the carboxy or amino terminus of the fusion gene These include known examples such as *NPEPPSP1-TBC1D3*, *NOTCH2NL-NBPF*, and *NPIP-PKD1P*^19,21,41^. We identify previously undescribed examples of such amalgamated duplicated genes, including *POLR2J-RASA4B* (Supplementary Fig. 12C), and *CYTOR-ANAPC1P4* (Supplementary Fig. 12D). We also identified a previously reported *LRRC37A2–NSFP1* fusion^37,40^; however, we find that this fusion appears to produce an ORF that extends from *NSFP1* into *LRRC37A2* exons (Extended Data Fig. 3E). *NSFP1* is a human-specific partial duplication derived from *NSF* and is located immediately adjacent to *LRRC37A2*. The two juxtaposed SDs span approximately 187.8 kbp and 75.6 kbp, respectively, for a combined length of 263.4 kbp, although we note that these potential gene innovations are not fixed in the human population.

### Protein-coding mutational constraint map

Previous analyses suggest duplicate genes acquiring a new function frequently become copy number invariant (e.g., *SRGAP2C*, *ARHGAP11B*) despite their increased potential for copy number variation via NAHR^14,17^. Unlike previous analyses^6,10^ that assessed copy number variation only at the gene-family level (Extend Data Fig. 4A), the pangenome enables copy number assessment for the most part at the individual paralog level (Extend Data Fig. 4B, Supplementary Table 2). Our study identified 755 SD genes (945 maximum copy counts) that are nearly fixed for copy number in the population, defined as genes retained as a single copy in more than 99% of haplotypes after excluding haplotypes affected by putative assembly errors (Fig. 1A). Of these, 276 (36.6%) represent ancestral genes, 302 (40.0%) are derived genes constrained with respect to copy number, and the remaining 177 (23.4%) are of undetermined origin. Predictably, some of the most copy number polymorphic duplicate genes correspond to the evolutionary youngest duplications or core duplicons^2^, which recurrently duplicate and exist at higher copy number in ape genomes (e.g., *SPDYE*, *NPIP*, *TBC1D3*). Nevertheless, even within highly variable gene families, we identify specific copies, such as *NPIPB2, NPIPB11,* and *NPIPB14P* (Supplementary Fig. 6C), and *NBPF8, NBPF9,* and *NBPF12* (Supplementary Fig. 10C) that are nearly fixed in copy number in the human population. The latter observation may be particularly relevant for identifying functional *NBPF* copies that are most likely involved in specifying basal radial glial cell identity in humans^42^.

**Fig. 4.**
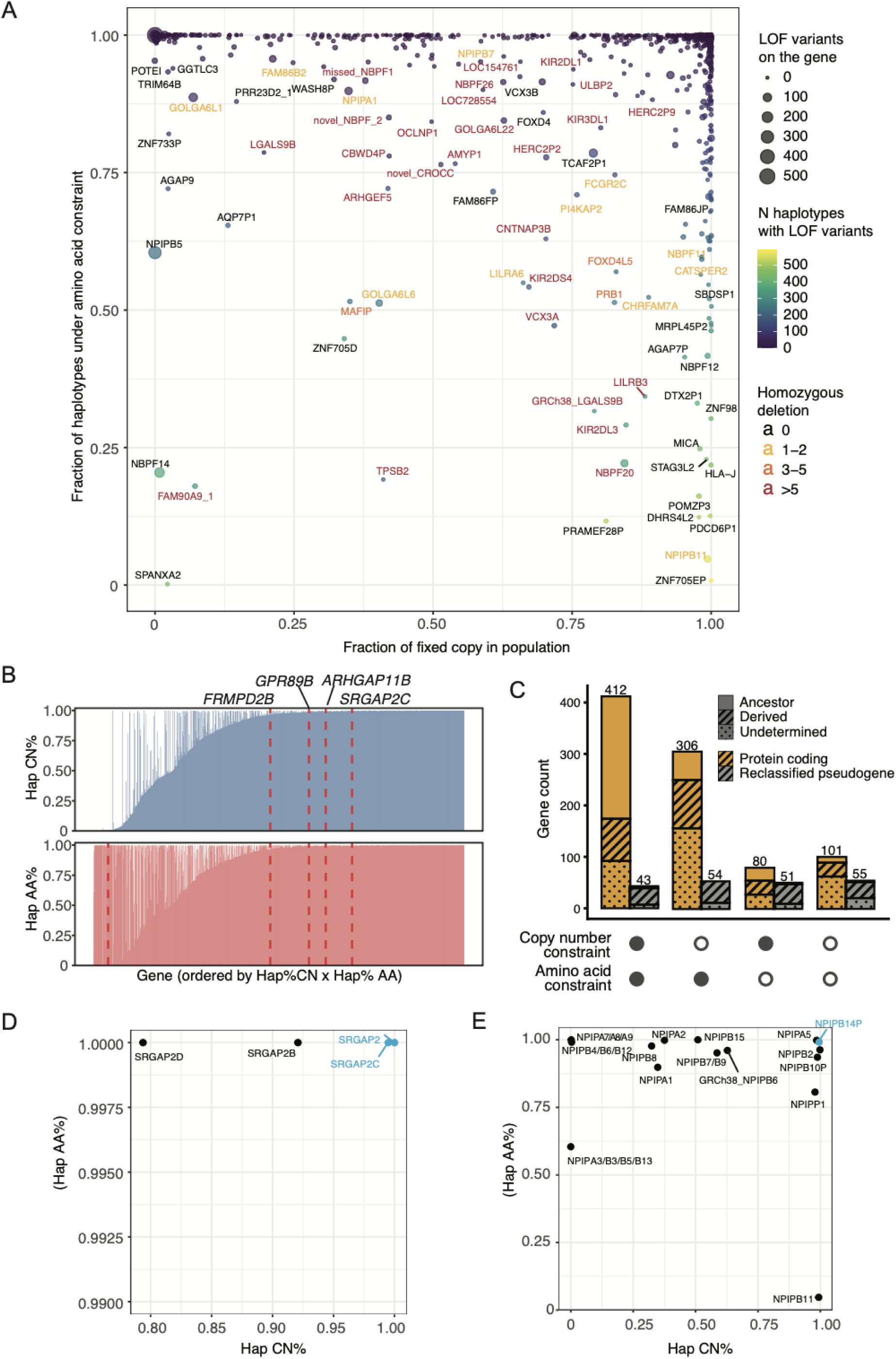
SD gene constraint. **(A)** Scatter plot of the constraint map for 1,102 monophyletic groups, showing the percentage of HPRC and HGSVC haplotypes (n = 596) carrying only one gene copy, after excluding assembly errors, versus the percentage of haplotypes carrying at least a gene copy under amino acid constraint. **(B)** Distribution of the percentage of haplotypes carrying only one gene copy (upper panel) and the percentage of haplotypes carrying at least one gene copy under amino acid constraint (lower panel). Dashed red lines highlight four known examples of functional derived genes involved in human brain development^12,14,43^. **(C)** Bar plot showing the number of paragroups identified from phylogenetic trees that were under copy number constraint, amino acid constraint, or both. Yellow indicates protein-coding paragroups, gray indicates reclassified pseudogene paragroups, and shading represents ancestral status, including ancestral, derived, and undetermined. Constraint scores for **(D)** *SRGAP2* (Supplementary Fig. 14) and **(E)** *NPIP* (Supplementary Fig. 6) gene families. Blue labeled genes are constrained at both copy number and amino acid levels. *NPIPB14P* is reclassified as a protein-coding gene.

Having assessed copy number constraint for individual paralogs based on phylogenetic subgrouping, we next evaluated amino acid constraint. Variants for each paralog were identified by comparison with the T2T-CHM13 reference gene sequence or, for the 42 non-CHM13 SD genes, with the most frequent sequences, and functional effects were predicted using the VEP^32^ to identify single-nucleotide variants (SNVs) predicted to cause deleterious changes in each putative protein-coding gene model. We define amino acid constrained paralogs as those harboring fewer predicted pathogenic loss-of-function (LOF) variants. Taken together, we conservatively define constrained SD genes as those in which more than 99% of haplotypes carry a single copy and exhibit amino acid constraint in more than 99% of haplotypes (Methods).

Returning to the *RGPD* gene family, *RGPD1*, *RGPD2*, and *RGPD8* are copy number stable across haplotypes, whereas the remaining paralogs exhibit substantial copy number variability. Both RGPD1 and RGPD2 show no detected nonsense variants (Fig. 2C, Supplementary Fig. 13). *RGPD2* is the one paralogous gene that is syntenic and orthologous among the African great apes, whereas *RGPD1* has been reported to be under positive selection^29^. Thus, both are under constraint. In contrast, *RGPD8*, despite being fixed in copy number, carries numerous predicted LOF variants. *RGPD3*, *RGPD4*, *RGPD5*, and *RGPD6* are all copy number variable and enriched for LOF variants (Fig. 2C, Supplementary Fig. 13).

We considered a total of 1877 genes, including 1188 protein-coding genes, 236 reclassified protein-coding genes, and 453 pseudogenes (Fig. 1C). Phylogenetic analysis grouped these genes into 1404 monophyletic groups, of which 1102 had gene models allowing constraint analysis, including 898 protein-coding groups and 204 reclassified groups. We then constructed a first-pass constraint map for these 1102 groups by jointly assessing population-level copy number fixation and amino acid constraint for each group (Fig. 4A-B, Supplementary Table 6). Following Iso-Seq-based gene model curation for 66 protein-coding genes, constraint classifications were further revised, with four genes changing their constraint status (*KRT81*, *ASAH2B*, *NOTCH2NLC* and *IL9R*) (Supplementary Figs. 12A, 15, 16). Overall, 24.2% of SD genes (455/1877) were classified as constrained at both the copy number and amino acid levels, including 412 protein-coding genes and 43 reclassified expressed pseudogenes (Fig. 4C). Of these 455 constrained SD genes, 242 were ancestral, 114 were derived, and 99 had undetermined ancestral status. Ancestral genes were constrained at a substantially higher rate than derived genes (64.5%, 242/375, versus 16.3%, 114/701). The constraint observed among this subset of derived genes suggests neofunctionalization. Among the constrained SD genes, 61 were human-specific duplications, including well-known examples such as *ARHGAP11B* and *SRGAP2C* (Fig. 4B,D). We also identified constrained copies within highly polymorphic core-duplication gene families, including *NPIB14P* (Fig. 4E), *SPDYE1/3/5/6* (Supplementary Fig. 7C), *LRRC37A3* (Supplementary Fig. 9C), and *NBPF9* (Supplementary Fig. 10C).

### Expression profiles of SD genes

Using the revised and novel gene models, we reassessed long-read transcriptomic data to provide a preliminary assessment of the expression landscape of each paralog (Methods). From an initial set of 5.6 billion reads, we excluded libraries with mixed tissue types, leaving 5.3 billion reads from 570 libraries across 83 tissues and 393 donors for analysis. As a control to assess tissue-specificity, we benchmarked against 515 tissue-specific unique genes identified from the Genotype-Tissue Expression (GTEx) Project. Highly expressed tissue-specific genes were well recovered in the Iso-Seq datasets (511/515), supporting the use of these data for duplicated gene expression profiling (Fig. 5A). Using criteria more stringent than those applied in the previous Iso-Seq gene model reassessment, defined as TPM > 5 in at least one Iso-Seq library or TPM > 1 in at least three libraries, we identified 1026 robustly expressed SD genes/pseudogenes. Next, we identified the top three tissues or cell types with the highest expression for each SD gene.

**Fig. 5.**
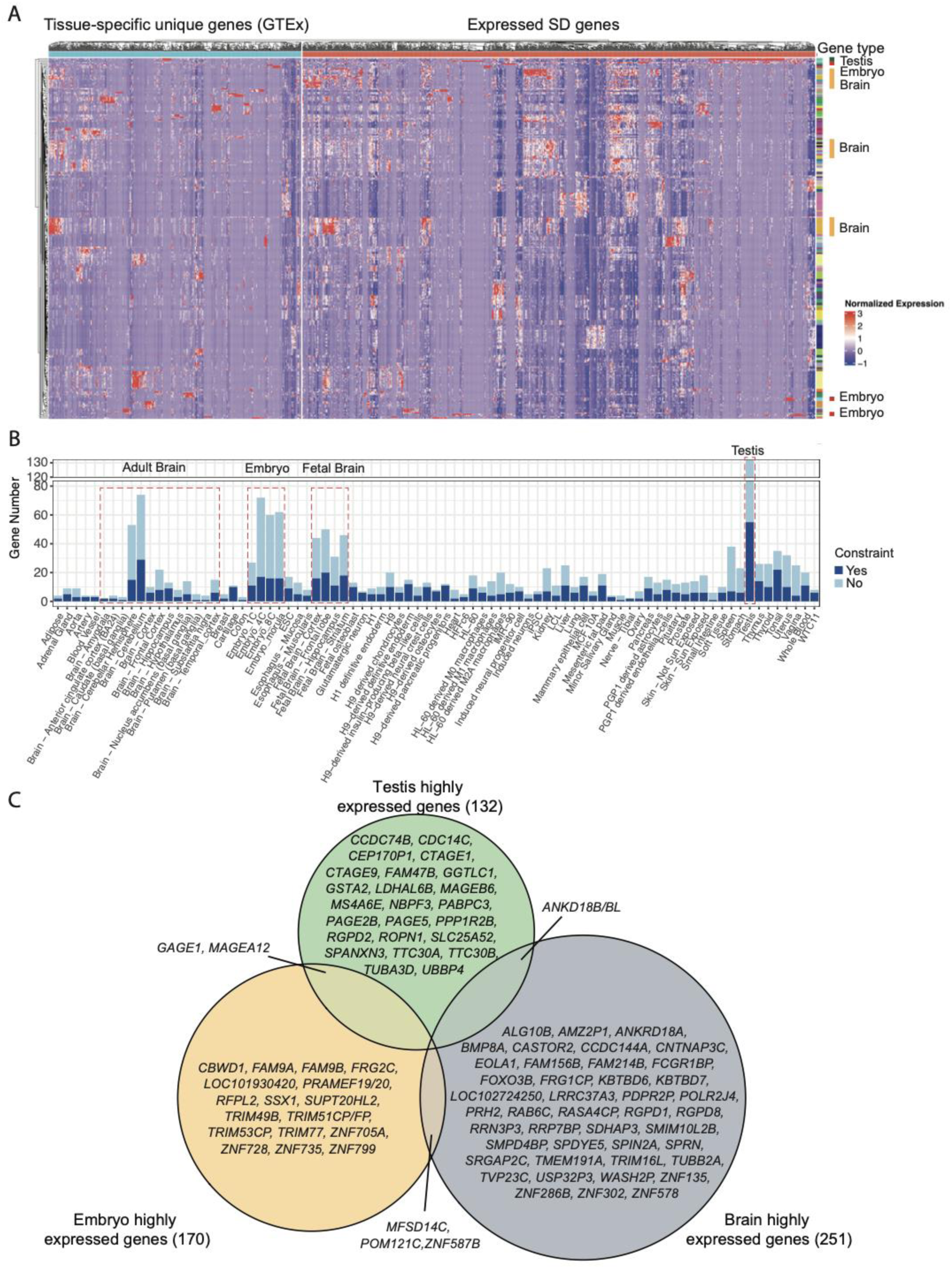
Gene expression of SD genes. **(A)** Heatmap of gene expression profiles from Iso-Seq data for tissue-specific genes identified from GTEx and expressed SD genes (TPM > 5 in at least one library or TPM > 1 in at least three libraries). **(B)** Highly expressed genes identified in each tissue or cell type. **(C)** Venn diagram showing the overlap among brain, embryo, and testis highly expressed SD genes. Genes shown are those derived or undetermined and under copy number and amino acid constraint.

We find that human SD genes are highly expressed in testis (132), embryo (170), and in the brain (251, representing the union of 132 adult-brain and 125 fetal-brain genes) (Fig. 5B, Supplementary Table 7). Compared with highly expressed unique genes, SD genes are significantly enriched in the testis (1.7-fold, p = 3.1 × 10⁻^7^) and early embryogenesis, including the 4-cell (2.3-fold, p = 9.5 × 10⁻⁸) and 8-cell embryo stage (2.0-fold, p = 8.7 × 10⁻⁵) (Supplementary Fig. 17). Direct GO enrichment analysis of 251 highly expressed brain and 170 embryonic SD genes revealed no significant functional enrichment, likely due to the limited functional annotation of SD genes. We therefore performed co-expression analysis between SD and unique genes, using the functional annotations of unique genes as a proxy to infer the biological roles of co-expressed SD genes (Methods). This approach identified 178 highly expressed brain SD genes co-expressed with highly expressed unique brain genes enriched in neuron-related pathways (Supplementary Fig. 18), and 159 highly expressed embryonic SD genes co-expressed with highly expressed unique embryonic genes involved in RNA processing and cell cycle/germ cell development pathways (Supplementary Fig. 19). Finally, we integrated expression profiles with copy number and amino acid constraint to prioritize the most likely functional SD genes. Among the 515 candidate genes highly expressed in brain, embryo, and testis, 185 showed evidence of constraint, including 92 ancestral genes, 56 derived genes, and 37 genes with undetermined status (Fig. 5C, Supplementary Table 7). Of the derived and undetermined constrained genes, 10 were human-specific, including the known *SRGAP2C*^12,17^ and *LRRC37A3*^44^. In addition, 25 of the constrained genes were reclassified expressed pseudogenes, including *AMZ2P1* and *FAHD2CP*.

## DISCUSSION

The availability of a human pangenome where high-quality linear references have been generated allowed us to assess a set of 1877 protein-coding genes/pseudogenes associated within SDs that have been difficult to interrogate since the initial sequencing of the human genome^45,46^. Analysis of the 458 recently released genomes with de novo gene annotations (Methods) also predicts another 621 non-reference SD gene families with as many as 878 genes (Fig. 1A). Previous studies estimated copy number variation of these duplicate genes based on hybridization intensity and read-depth analysis of short-read sequencing^5,6,10,47,48^. These studies often assessed variation in the aggregate (i.e., at the gene-family level) as opposed to being paralog-specific. Other LRS-based methods have been developed to distinguish paralogous variants within duplicated gene families, but these approaches largely focused on a limited set of well-known, disease-associated loci^49^ or a modest number of genomes^26,37^. To help mitigate the high sequence identity of many of these loci due to either recent duplication or IGC, we applied a population-based phylogenetic approach to systematically classify all genes into groups—where the frequency of each paralog or paragroup could be assessed as well as SNVs that might alter amino acid composition. Coupling these phased genomes with long-read transcriptome data then allowed gene models to be revisited by mapping full-length transcripts to specific loci helping to alleviate some of the challenges associated with assigning transcript data and subsequent gene annotation of these regions^50,51^.

Among the 1877 candidates, we identified 1126 protein-coding gene candidates with evidence of expression, supported by at least five Iso-Seq reads for an isoform containing a complete ORF. Within this expressed set, 150 gene models diverged from current reference annotations, and 236, corresponding to 35.1% of the 672 annotated pseudogenes, were reclassified as potentially transcribed protein-coding genes with well-defined ORFs, a distinction made possible by the paralog-specific resolution of long-read sequencing. A large number of these revised models arise as a result of a juxtaposition of SDs of diverse evolutionary origins (Extend Data Fig. 3E, Supplementary Figs. 12C-D). While the biological significance of many of these “fusion genes” requires further follow-up, these features are reminiscent of the exon shuffling model put forward by Walter Gilbert in 1978^52^. Application of both Fiber-seq data and mass spectrometry data for a subset of these provided further support of promoter activity and protein potential, respectively. Among these are several human-specific genes now associated with brain development^17,53^ that were originally classified as pseudogenes until careful transcript definition followed by experimental functional characterization proved otherwise^12,15,16,54,55^. In other cases, the fusions involve genes with co-evolved functions, such as *NBPF14* and *NOTCH2NLB*, where *NBPF* genes have been shown to promote basal progenitor formation through oblique cleavage, while *NOTCH2NLB* expands the apical progenitor pool, jointly facilitating brain evolution^56^. We suggest that this multiomic pangenomic approach we developed can be more generally applied to other species to quickly distinguish likely functional protein-coding genes from true pseudogenes and to assess their relative frequency at the population level.

We note that we did not find sufficient Iso-Seq support (Methods) for 749 RefSeq gene models. In other cases, long-read transcript data did not predict a complete ORF model (n=68 cases). This lack of transcript support may reflect the limited range of human tissues surveyed or technical challenges in recovering full-length transcripts by Iso-Seq. For example, *OPN1LW* and *OPN1MW* are well-known opsin genes required for the detection of the red-green wavelength spectrum^57^, but whose expression is largely localized to the outer segments of cone photoreceptor cells. This tissue was not adequately sampled as part of our long-read transcriptome database. In other cases, exceptionally long or repeat-rich coding regions hinder complete isoform recovery (n=36). The large *HYDIN2* pseudo/gene has an ORF and an expected transcript length of 10.5 kbp^53^, but we could not identify complete Iso-Seq isoforms for this locus. Similarly, *NBPF10/14/19* (10.5 kbp) contain massive expansions of the human-specific Olduvai domain repeat expansions^58^ that were not fully recovered in the transcript data.

It has long been recognized that recent duplicate genes provide the substrate for the emergence of new genes and can follow a variety of different evolutionary fates, ranging from redundant function, pseudogenization, subfunctionalization to neofunctionalization^59–61^. To investigate the transcriptomic fates of derived paralogs, we compared tissue-expression patterns between ancestral genes and their derived copies across 697 ancestral-derived gene pairs from 375 gene families (Supplementary Table 8). The most frequent is asymmetry expression, defined as ancestral- or derived-copy dominance in which one copy shows substantially higher expression than the other across all expressed tissues (375 pairs; fold change > 2). This was followed divergent expression, in which the derived copy showed either a different tissue of maximum expression or at least a twofold tissue-specific increase relative to the ancestral copy (133 pairs); partially divergent expression, in which the derived copy was expressed in only a subset of the tissues in which the ancestral copy was expressed (105 pairs); and dosage-balanced expression, in which both copies are similarly expressed across tissues (24 pairs; correlation >0.9 and fold change <2). In the remaining 60 pairs, neither the ancestral nor the derived copy was found to be expressed.

Among the 133 derived genes with divergent expression relative to their ancestral copies and the 15 genes showing derived-copy dominance, 78 exhibited a shift toward adult brain-, fetal brain-, or embryo-specific expression, or showed increased expression in one or more of these tissues. Of these 78 genes, 26 were under constraint. These expression shifts suggest regulatory divergence following duplication and identify candidate genes that may have acquired new tissue-specific functions, representing potential neofunctionalization candidates for further investigation. Examples include the derived genes *CBWD1* and *POM121C*, both of which have been reported to be associated with intellectual developmental disorders^62,63^. Compared with their ancestral copies, both genes showed high embryo-specific expression and were under both copy number and amino acid constraint (Supplementary Fig. 20).

Because typically thousands to tens of thousands of genomes are required to statistically define mutational constraint^23^, we regard our joint analysis of amino acid and copy number constraint as a first pass because it was defined on less than 600 haplotype-resolved genomes. Nevertheless, the approach clearly distinguishes the most copy number polymorphic gene families in the human genome (*DUX*, *GAGE12J*, *TBC1D3*, etc.) and, therefore, gene families that would benefit the most from the pangenome project for improved genotyping and association. It also identifies specific loci within copy number variant gene families that are relatively most constrained including those already associated with function (*SRGAP2C*^12^, *ARHGAP11B*^14^, *CROCCP2*^15^, etc.) but a larger subset that would benefit the most from targeted functional analyses in model organisms^43^.

A limitation of our strategy, however, is that it is distinctly monogenic and haplotype biased, not accounting for the joint contribution of highly similar genes with likely overlapping function within an individual. For example, duplicated genes such as *SMN1*, whose deletion results in spinal muscular atrophy (SMA), and *SMN2* are both polymorphic. Because *SMN2* can compensate, at least in part, for the loss of *SMN1*, SMA severity depends on the total copy number of both genes^64^. If we relax the copy number and amino acid constraint criterion from the haplotype level to the diplotype level, we identify 221 such genes that are constrained at the diplotype level in the human population (Supplementary Table 6). Although these genes show substantial haplotype-level copy number polymorphism, each individual retains at least one copy. In addition to several well-known highly copy number variable genes, including *AMY2A*^65^, *C4A*^66^, *FCGR3B*^67^, and *TPSAB1*^68^, we also identified *GPRIN2*, reported associated with neuronal projection^69,70^, as well as several paired duplicates potentially affected by gene conversion, such as *NOTCH2NLA-NOTCH2NLB*, *GTF2IRD2*–*GTF2IRD2B* and *TRIM43*–*TRIM43B*, which may function together.

Several other challenges remain. First, our analysis used T2T-CHM13 as the starting reference. Although population-scale assemblies allowed us to recover some genes missing from T2T-CHM13, gene families absent from T2T-CHM13 still exist and we estimate 621 additional non-reference, lower frequency SD gene families that will require follow-up (Table 1). Related to this, the most identical duplicate genes, in general, are highly copy number polymorphic, but the Iso-Seq datasets used here are not matched to the same donor-specific genome assemblies. As a result, when a duplicated gene is not detected in a transcriptome library, it can be difficult to distinguish an absence of expression from absence of that gene copy in the donor genome. Future studies using donor-matched genome assemblies where long-read transcriptomes and regulatory datasets have been produced across multiple tissues, such as the Somatic Mosaicism across Human Tissues (SMaHT) Network^71^, will be important for systematically characterizing the regulation and function of duplicated genes. Finally, now that 386 new protein models and 455 constrained genes have been identified, more directed proteomic approaches can be designed to identify unique peptides by mass spectra that distinguish closely related peptides^72,73^. Similarly, targeted population genetic studies formally assessing different models of selection can be performed on larger, population-level LRS datasets being developed as part of biobanks such as the *All of Us* cohort^74^. Discovering gene-disruptive mutations in these constrained or adaptive genes that result in phenotypes may be possible given that such datasets are associated with extensive electronic health record data^75^.

## METHODS

### Duplicated gene families identification

Using the T2T-CHM13 v2.0 reference genome, we selected genes longer than 1 kbp, including both protein-coding genes and pseudogenes, in which at least one paralog within the gene family had at least 30% of its gene body overlapping SD regions. We retained only gene families in which at least one member was annotated as protein-coding. We then performed all-to-all sequence alignments among these genes using BLAST^76^ (v2.12.0). Genes were grouped into the same family based on reciprocal overlap criteria, requiring each alignment to cover at least 70% of one gene and one-third of the other, with a minimum sequence identity of 90%. Gene families were excluded from further analysis if they were located on sex chromosome pseudoautosomal regions and contained one member on chromosome X and another on chromosome Y.

### Duplication time and ancestor identification

For each gene family, gene sequences were extracted with flanking regions ranging from 0 bp to 20 kbp, based on the distance between paralogs within the family. Orthologous sequences were retrieved using minimap2^77^ (v2.28) from ten nonhuman primate species: chimpanzee (*Pan troglodytes*, PTR), bonobo (*Pan paniscus*, PPA), gorilla (*Gorilla gorilla*, GGO), Bornean orangutan (*Pongo pygmaeus*, PPY), Sumatran orangutan (*Pongo abelii*, PAB), siamang (*Symphalangus syndactylus*, SSY), rhesus macaque (*Macaca fascicularis*, MFA), common marmoset (*Callithrix jacchus*, CJA), olive baboon (*Papio anubis*, ANA), and slow loris (*Nycticebus coucang*, LCT). Sequences were aligned using MAFFT^78^ (v7.525), and phylogenetic trees were reconstructed using IQ-TREE^79^ (v2.1.2), with the most distantly related species serving as the outgroup. Duplication time was estimated from the most recent common ancestor (MRCA) of the human duplicated loci in the gene tree. When tree topology indicated that a duplication predated the human lineage, we identified ancestral copies using sequences from species that diverged before the MRCA node and retained only a single copy of the gene.

### Phylogenetic trees construction and paralog grouping

For each gene family, we extracted gene sequences from HPRC and HGSVC human assemblies, two reference genomes (T2T-CHM13 and GRCh38), and nonhuman primate assemblies, including PTR, PPA, GGO, PPY, PAB, SSY, and MFA. Gene sequence extraction was performed using minimap2^77^ (v2.28) with the following parameters (-x asm20 -c –secondary=yes -p 0.3 -N 10000 –eqx -r 500 -K 500M). Nonhuman primate genes were used as outgroups. Multiple sequence alignments (MSAs) were generated using MAFFT^78^ (v7.525). We used intronic regions to build phylogenetic trees with IQ-TREE^79^ (v2.1.2); for genes lacking intronic regions or with <500 bp of intronic sequence or with high sequence identity (>99%), the full gene sequence was used for tree construction. Tips with abnormally long branch length and partial gene duplications were removed prior to downstream analyses. Paralogs were defined on the basis of phylogenetic clustering. Genes forming clades with bootstrap support >90 were assigned to the same paralog group. Clades with bootstrap <90 were reevaluated and reassigned. We identified clades showing intra-group variation up to 1.5 times the allelic variation observed in SD regions, corresponding to 15.3 SNVs per 10 kbp^4^. A population-level paralog group was required to constrain at least 100 independent paralogs.

### Gene conversion identification

We split each gene sequence from assemblies into 500 bp bins using a 200 bp sliding step. Each window was then aligned back to the T2T-CHM13 v2.0 reference genome using minimap2^77^ (v2.28). The best match for each window was defined as the alignment with the highest sequence identity. If a window mapped to gene conversion pairs with identical sequence identity, it was classified as an equal match.

### Gene model and ORF prediction

Iso-Seq reads were aligned to the T2T-CHM13 v2.0 reference genome using pbmm2 (v1.17.0, https://github.com/PacificBiosciences/pbmm2) with the ISOSEQ preset. Uniquely mapped reads were retained for downstream analysis, and secondary alignments were removed. For each gene, we extracted sub BAM files containing reads within the gene region from a total of 583 libraries (5.6 billion reads across 83 tissue/cell types) and merged these sub BAM files for downstream analysis. Redundant transcripts were collapsed into unique isoforms based on exonic structure using Iso-Seq collapse. Transcripts were then classified using PacBio Pigeon (v1.2.0, https://isoseq.how/classification/pigeon.html), and low-quality isoforms were removed using Pigeon filter. ORFs were annotated using TransDecoder as implemented in SQANTI3^80^ (v5.2). For each gene, isoforms shorter than 50% of the gene length were excluded, and the most abundant remaining gene model was selected and compared against RefSeq annotations. Transcript structures were visualized using pyGenomeTracks^81^ (v3.9).

### Variant detection and annotation

Gene sequences were aligned using an MSA with MAFFT^78^ (v7.525). SNVs and indels were identified by comparing each gene sequence to the corresponding reference gene sequence from CHM13 v2.0. Variant functional consequences were annotated using VEP^32^ (v111) with gene annotation (JHU RefSeqv110 + Liftoff v5.2). For potentially expressed pseudogenes or genes with alternative models inferred from Iso-Seq data rather than RefSeq, gene models were obtained from Iso-Seq predictions. For variant detection and annotation of novel genes absent from both T2T-CHM13 and GRCh38, we first selected a representative assembly sequence for each gene from the MSA. The representative sequence was defined as the assembly sequence with the smallest distance to the clade consensus, thereby capturing the most common sequence pattern observed across assemblies. We then mapped Iso-Seq reads to the representative assembly sequence to identify gene models using SQANTI3^80^ (v5.2). Gene models with predicted ORFs and support from more than five reads were retained, and their Gene Transfer Format (GTF) annotations were generated for downstream analyses. Variants were then called against the representative assembly sequence, and their functional consequences were annotated using VEP^32^. Pathogenicity predictions for missense variants were obtained using AlphaMissense^82^, which were lifted over to T2T-CHM13 coordinates using UCSC LiftOver and complemented by PolyPhen and SIFT (20240502). Lollipop plots were generated using the R package trackViewer^83^ (v1.40.0).

### Constraint analysis

SD gene constraint was assessed at both the copy number and amino acid levels for each paralog clade identified from the phylogenetic tree, using two scores. The copy number score was calculated as the percentage of haplotypes carrying only a single copy of the gene, across all haplotypes for that paralog after excluding those with copies located in regions of assembly error. The amino acid score was calculated using this same set of assembly-error-filtered haplotypes, as the percentage carrying at least one copy free of nonsense variants. A haplotype with multiple copies was counted as constrained so long as at least one of its copies lacked a nonsense variant. To identify copy-number-polymorphic genes that are nonetheless under constraint, we evaluated genes that did not meet haplotype-level constraint criteria at the diploid level instead. Specifically, we calculated (i) the percentage of individuals in which at least one of the two haplotypes carried at least one gene copy, and (ii) the percentage of individuals in which at least one of the two haplotypes carried at least one copy free of nonsense variants. We defined paralogs as constrained when both the copy number score and the amino acid score exceeded 0.99 (i.e., >99%).

### Pseudogene accessibility analysis

Revived pseudogene loci, dead pseudogene loci, and a protein-coding positive control were extracted from HPRC assemblies and HG002 for the 78 haplotypes with paired Fiber-seq data^25^. For each gene, a TSS window was defined as ±500 bp around the TSS. Per-haplotype Fiber-seq peak calls generated by Minkina, Mah-Som et al. (in preparation) were intersected with the TSS windows to obtain peak presence per locus and maximum Fiber-seq actuation per locus. To control for differences in set size, dead and protein-coding loci were each randomly subsampled to match the number of revived pseudogene loci. Pairwise enrichment between the three classes was assessed with two-sided Fisher’s exact tests on 2×2 tables of (locus has peak) × (class).

### Public proteomics mass spectrometry data processing

Sequences of queried proteins were added to the FASTA file before sequences of annotated human proteome from Uniprot (12/2025) containing Swiss-Prot canonical and isoform sequences. Common contaminants and decoys for all proteins were included at the beginning and end, respectively. Raw files were searched against the final FASTA file. Comet search algorithm was used to match peptides to spectra, with parameters depending on the type of analysis. Parameters for label-free DDA from LTQ-Orbitrap Velos/Elite mass spectrometer: precursor mass tolerance: 20 ppm, fragment mass tolerance: 0.05 Da. Parameters for TMT-DDA (10plex or 11plex) from Orbitrap Fusion Lumos or Q Exactive HF-X mass spectrometers: MS2: precursor mass tolerance: 20 ppm, fragment mass tolerance: 0.02 Da; MS3: precursor mass tolerance: 20 ppm, fragment mass tolerance: 1 Da, reporter ion tolerance: 0.003 Da, MS2 isolation width: 0.7, MS2 isolation width: 1.2. Peptide static modification included carbamidomethyl (57.0214637236) on cysteine residues for all datasets, and TMT (229.162932) on lysine residues and peptide N-termini. Variable modification included oxidation (15.9949146221) on methionine residues. Peptides with up to two missed cleavages were included. Peptide-spectrum matches (PSMs) were filtered to a 2% false discovery rate (FDR) using a linear discriminant analysis. Proteins were filtered to a 2% FDR using the protein picker method and assembled following protein parsimony 12. Peptides in TMT-labeled samples analyzed with MS3 were required to have a summed TMT reporter ion signal-to-noise (SnSum) ≥100. Peptides that matched to queried proteins, including pseudogenes, were analyzed and compared with other proteins in the data for their quality using the peptide metrics.

### Gene expression analysis

Updated gene models, including reclassified pseudogenes, gene fusions, non-CHM13 gene models, and protein-coding genes with revised gene structures derived from Iso-Seq data, were incorporated into the reference annotation GTF file. Non-CHM13 gene sequences were added to the T2T-CHM13 reference genomes. Gene-level expression was quantified from long-read transcriptomic data using IsoQuant^84^ (v3.10.0). Genes were retained for downstream analysis if they had TPM >5 in at least one library or TPM >1 in at least three libraries. Tissue-specific genes were identified from the GTEx short-read dataset for quality control, with genes considered tissue-specific if they showed a log2 fold change >2 in one tissue compared to all others. To identify highly expressed SD genes within each tissue, tissues were ranked by expression level for each gene and the top three ranked tissues were evaluated. If the top-ranked tissue had a TPM value at least 50 higher than the second-ranked tissue, only the top-ranked tissue was retained. Similarly, if the second-ranked tissue had a TPM value at least 50 higher than the third-ranked tissue, the third-ranked tissue was excluded. For each tissue type, expression profiles of both unique and SD highly expressed genes were used as input for weighted gene co-expression network analysis (WGCNA)^85^. Co-expression modules were identified, and genes with a module membership coefficient (kME) >0.5 were retained. This filtering step removed genes that were only weakly connected to their assigned module, ensuring that the retained unique and duplicated genes closely reflected the co-expression pattern representative of that module. For each module, unique genes were used for Gene Ontology (GO) enrichment analysis, with a significance threshold Benjamini-Hochberg adjusted p-value < 0.05, to infer the biological functions of co-expressed duplicated genes.

### Detection of non-reference SD genes

We used 458 HPRC2 assemblies that contain both gene annotation as well as short-read validated SD tracks. Based on the gene annotation (https://github.com/human-pangenomics/hprc_intermediate_assembly/blob/main/data_tables/annotation/cat/cat_genes_hprc_r2_v1.3.index.csv) available for each of the HPRC2 assemblies, we defined SD genes, under the following criteria: (1) the gene body overlapping with at least 30% with SD track, (2) the overlap region needs to include at least one exon, (3) considered genes contain at least two exons and minimum length of 1 kbp, and (4) the gene biotype of the protein-coding gene was considered. The reference annotation (JHU RefSeqv110 + Liftoff v5.2) was compared with the maximum paralog gene counts observed across the pangenome. We classified genes to ‘non-reference SD gene’ or ‘copy number increase’ if higher copy number was observed in the pangenome compared to the reference, and remaining genes which show equal number of gene counts were classified into ‘reference SD gene’.

## DATA AND CODE AVAILABILITY

All HPRC data, including assemblies and PacBio Kinnex reads, are available at: https://data.humanpangenome.org/raw-sequencing-data. Fiber-seq data are available at https://s3-us-west-2.amazonaws.com/humanpangenomics/index.html?prefix=submissions/5ECA1D3E-1C37-44B7-BA20-576417F786F0-UCSC_HPRC_ONT_YEAR1_FIBERSEQ/.

All HGSVC assemblies are available at: https://ftp.1000genomes.ebi.ac.uk/vol1/ftp/data_collections/HGSVC3/release.

Iso-Seq data are available at the accessions listed in Supplementary Table 3.

All mass spectrometry proteomics data were downloaded from the Proteomics data commons (https://pdc.cancer.gov) or PRIDE databases. Datasets included: human proteome map GTEx (PXD016999), human adult and fetal proteome (PXD000561), Clinical Proteomic Tumor Analysis Consortium (CPTAC) LUAD (PDC000489), CPTAC PNET (PDC000590), CPTAC GBM (PDC000204), Broad Institute Medulloblastoma (PDC000433), CPTAC HNSCC (PDC000221), and Proteogenomic Translational Research Center (PTRC) AML (PDC000477).

Code for building phylogenetic trees and paralog-level variant identification and classification is available at https://github.com/LuyaoRen/pangenome-sd-genes-workflows. All other code is publicly available.

## ACKNOWLEDGMENTS

We thank Tonia Brown for editing the manuscript and supplementary materials. We would like to acknowledge the U.S. National Human Genome Research Institute (NHGRI) of the National Institutes of Health (NIH) for funding the following grants supporting the creation of the human pangenome reference: U41HG010972, U01HG010971, U01HG013760, U01HG013755, U01HG013748, U01HG013744, R01HG011274, and the Human Pangenome Reference Consortium (BioProject ID: PRJNA730823). Research reported in this publication was supported, in part, by the NHGRI under Award Numbers U24HG007497 and R01HG002385 to E.E.E. This research was supported in part by the Intramural Research Program of the NIH. M.R.V. was supported by a Pathway to Independence award from the National Institute of General Medical Sciences (4R00GM155552). D.K.S. acknowledges support from the Pew Charitable Trusts. The content is solely the responsibility of the authors and does not necessarily represent the official views of the NIH. E.E.E. is an investigator of the Howard Hughes Medical Institute.

## COMPETING INTERESTS

E.E.E. is a scientific advisory board (SAB) member of Variant Bio, Inc. D.K.S. is a collaborator with ThermoFisher Scientific. All other authors declare no competing interests.

## AUTHOR CONTRIBUTIONS

E.E.E. and L.R. conceptualized the study. K.H. and K.M.M. generated the data. L.R., D.Y., K.V., M.R.V., P.C.D., X.G., Y.K., and J.L. conducted formal analyses and created visualizations. L.R. and E.E.E. did the interpretation. L.R. and E.E.E wrote the original draft. E.E.E., D.K.S., M.R.V., and A.B.S. supervised the study. All authors reviewed and edited the manuscript.

## Human Pangenome Reference Consortium Version 2 Authors

Derek Albracht^1^, Ivan A. Alexandrov^2^, Jamie Allen^3^, Alawi A. Alsheikh-Ali^4^, Nicolas Altemose^5^, Casey Andrews^6^, Dmitry Antipov^7^, Lucinda Antonacci-Fulton^1^, Alexander Arguello^8^, Mobin Asri^9^, Marcelo Ayllon^10^, Jennifer R. Balacco^11^, Floris P. Barthel^12^, Edward A. Belter Jr^1^, Halle D. Bender^9^, Andrew P. Blair^9^, Davide Bolognini^13^, Katherine E. Bonini^14^, Christina Boucher^15^, Guillaume Bourque^16,17,18^, Silvia Buonaiuto^19^, Shuo Cao^19^, Andrew Carroll^20^, Ann M. Mc Cartney^21^, Monika Cechova^9^, Mark J.P. Chaisson^22^, Pi-Chuan Chang^20^, Xian Chang^9^, Jitender Cheema^3^, Haoyu Cheng^23^, Claudio Ciofi^24^, Hiram Clawson^9^, Sarah Cody^1^, Vincenza Colonna^19^, Holland C. Conwell^25^, Robert Cook-Deegan^26^, Mark Diekhans^9^, Maria Angela Diroma^24^, Daniel Doerr^27,28,29^, Zheng Dong^6^, Danilo Dubocanin^5^, Richard Durbin^30,31^, Jana Ebler^27,32^, Evan E. Eichler^10,33^, Jordan M. Eizenga^9^, Parsa Eskandar^9^, Eddie Ferro^15^, Anna-Sophie Fiston-Lavier^34,35^, Sarah M. Ford^25^, Willard W. Ford^36^, Giulio Formenti^11^, Adam Frankish^3^, Mallory A. Freeberg^3^, Qichen Fu^6^, Stephanie M. Fullerton^37^, Robert S. Fulton^1^, Shenghan Gao^38^, Yan Gao^39^, Gage H. Garcia^10^, Obed A. Garcia^40^, Joshua M.V. Gardner^9^, Shilpa Garg^41^, Erik Garrison^19^, Nanibaa’ A. Garrison^42,43,44^, John E. Garza^1^, Margarita Geleta^45,46^, Mohammadmersad Ghorbani^47^, Tina A. Graves-Lindsay^1^, Richard E. Green^25^, Carol W. Greider^48^, Cristian Groza^49^, Bida Gu^22^, Andrea Guarracino^12,19^, Melissa Gymrek^50^, Maximilian Haeussler^9^, Leanne Haggerty^3^, Ira M. Hall^51,52^, Nancy F. Hansen^7^, Yue Hao^12^, Mohammad Amiruddin Hashmi^4^, David Haussler^9^, Prajna Hebbar^9^, Peter Heringer^27,28,29^, Glenn Hickey^9^, Todd L. Hillaker^9^, S. Nakib Hossain^3^, Neng Huang^39,53^, Sarah E. Hunt^3^, Toby Hunt^3^, Alexander G. Ioannidis^5,9,46^, Nafiseh Jafarzadeh^9^, Nivesh Jain^11^, Erich D. Jarvis^11,33^, Maryam Jehangir^12^, Juan Jiang^6^, Eimear E. Kenny^14^, Juhyun Kim^7^, Bonhwang Koo^11^, Sergey Koren^7^, Milinn Kremitzki^1,6^, Charles H. Langley^54^, Ben Langmead^55^, Heather A. Lawson^6^, Daofeng Li^6^, Heng Li^39,53^, Ronghan Li^6^, Wen-Wei Liao^51,52^, Jiadong Lin^10^, Tianjie Liu^6^, Glennis A. Logsdon^38^, Ryan Lorig-Roach^9^, Jonathan LoTempio Jr^21,56^, Hailey Loucks^9^, Jane E. Loveland^3^, Jianguo Lu^57^, Shuangjia Lu^51,52^, Julian K. Lucas^9^, Walfred Ma^22^, Juan F. Macias-Velasco^1,6,58^, Kateryna D. Makova^59^, Maximillian G. Marin^39,53^, Christopher Markovic^1^, Tobias Marschall^27,32^, Franco L. Marsico^19^, Fergal J. Martin^3^, Mira Mastoras^9^, Capucine Mayoud^34^, Brandy McNulty^9^, Jack A. Medico^11^, Julian M. Menendez^9^, Karen H. Miga^9^, Anna Minkina^60^, Matthew W. Mitchell^61^, Saswat K. Mohanty^62^, Younes Mokrab^47,63,64^, Jean Monlong^65^, Shabir Moosa^47^, Avelina Moreno-Ochando^66,67^, Shinichi Morishita^68^, Jonathan M. Mudge^3^, Katherine M. Munson^10^, Njagi Mwaniki^69^, Nasna Nassir^4^, Chiara Natali^24^, Shloka Negi^9^, Lingbin Ni^10^, Adam M. Novak^9^, Faith Okamoto^9^, Keisuke K. Oshima^38^, Pilar N. Ossorio^70,71^, Chie Owa^68^, Sadye Paez^11^, Benedict Paten^9^, Clelia Peano^13,72^, Adam M. Phillippy^7,55,73,74^, Brandon D. Pickett^7^, Laura Pignata^19^, Nadia Pisanti^69^, David Porubsky^10,75^, Pjotr Prins^19^, Timofey Prodanov^27,32^, Anandi Radhakrishnan^9^, T. Rhyker Ranallo-Benavidez^12^, Brian J. Raney^9^, Mikko Rautiainen^76^, Alessandro Raveane^13^, Andreas Rechtsteiner^48^, Luyao Ren^10,33^, Arang Rhie^7^, Fedor Ryabov^77,78^, Samuel Sacco^25^, Farnaz Salehi^19^, Michael C. Schatz^55,79^, Laura B. Scheinfeldt^80^, Aarushi Sehgal^36^, William E. Seligmann^25^, Mahsa Shabani^81^, Kishwar Shafin^20^, Shadi Shahatit^34^, Ruhollah Shemirani^14^, Vikram S. Shivakumar^55^, Swati Sinha^3^, Jouni Sirén^9^, Linnéa Smeds^62^, Steven J. Solar^7^, Marco Sollitto^11,24^, Nicole Soranzo^13,30,82^, Andrew B. Stergachis^10,60^, Marie-Marthe Suner^3^, Yoshihiko Suzuki^68^, Arda Söylev^27,32^, Ahmad Abou Tayoun^83,84^, Jack A.S. Tierney^3^, Chad Tomlinson^1^, Francesca Floriana Tricomi^3^, Mohammed Uddin^4,85^, Matteo Tommaso Ungaro^25,86^, Rahul Varki^15^, Flavia Villani^19^, Ivo Violich^9^, Mitchell R. Vollger^87^, Brian P. Walenz^7^, Charles Wang^88^, Lisa E. Wang^14^, Ting Wang^1,6,58^, Aaron M. Wenger^89^, Conor V. Whelan^11^, Zilan Xin^6^, Zheng Xu^6^, Kai Ye^90^, DongAhn Yoo^10^, Wenjin Zhang^6^, Ying Zhou^39^, Xiaoyu Zhuo^6^, Giulia Zunino^13^

### Affiliations

^1^ McDonnell Genome Institute, Washington University School of Medicine, St. Louis, MO 63108, USA

^2^ Department of Human Molecular Genetics and Biochemistry, Faculty of Medical and Health Sciences, Tel Aviv University, Tel Aviv 69978, Israel

^3^ European Molecular Biology Laboratory, European Bioinformatics Institute (EMBL-EBI), Wellcome Genome Campus, Hinxton, Cambridge CB10 1SD, UK

^4^ Center for Applied and Translational Genomics (CATG), Mohammed Bin Rashid University of Medicine and Health Sciences, Dubai Health, Dubai, UAE

^5^ Department of Genetics, Stanford University, Palo Alto, CA 94304 USA

^6^ Department of Genetics, Washington University School of Medicine, St. Louis, MO 63110, USA

^7^ Genome Informatics Section, Center for Genomics and Data Science Research, National Human Genome Research Institute, National Institutes of Health, Bethesda, MD 20892, USA ^8^ Division of Genome Sciences, National Human Genome Research Institute, Bethesda, MD 20871 USA

^9^ UC Santa Cruz Genomics Institute, University of California, Santa Cruz, CA 95060, USA

^10^ Department of Genome Sciences, University of Washington School of Medicine, Seattle, WA 98195, USA

^11^ The Vertebrate Genome Laboratory, The Rockefeller University, New York, NY 10065, USA

^12^ Bioinnovation and Genome Sciences, The Translational Genomics Research Institute (TGen), Phoenix, AZ 85004, USA

^13^ Human Technopole, Milan, Italy

^14^ Institute for Genomic Health, Icahn School of Medicine at Mount Sinai, New York, NY 10029, USA

^15^ Department of Computer and Information Science and Engineering, University of Florida, Gainesville, FL 32611, USA

^16^ Canadian Center for Computational Genomics, McGill University, Montréal, QC H3A 0G1, Canada

^17^ Department of Human Genetics, McGill University, Montréal, QC H3A 0G1, Canada

^18^ Victor Phillip Dahdaleh Institute of Genomic Medicine, Montréal, QC H3A 0G1, Canada

^19^ Department of Genetics, Genomics and Informatics, University of Tennessee Health Science Center, Memphis, TN 38163, USA

^20^ Google LLC, Mountain View, CA 94043, USA

^21^ Institute of Clinical and Translational Sciences, University of California, Irvine, CA 92697, USA

^22^ Quantitative and Computational Biology, University of Southern California, Los Angeles, CA 90089, USA

^23^ Department of Biomedical Informatics and Data Science, Yale School of Medicine, New Haven, CT 06510, USA

^24^ Department of Biology, University of Florence, Sesto Fiorentino, FI 50019, Italy

^25^ Department of Ecology and Evolutionary Biology, University of California, Santa Cruz, CA 95060, USA

^26^ Arizona State University, Consortium for Science, Policy & Outcomes, Washington, DC 20006, USA

^27^ Center for Digital Medicine, Heinrich Heine University Düsseldorf, Düsseldorf, NRW, DE

^28^ Department for Endocrinology and Diabetology at the Medical Faculty and University Hospital Düsseldorf, Heinrich Heine University Düsseldorf, Düsseldorf, NRW, DE

^29^ Paul-Langerhans-Group Computational Diabetology, German Diabetes Center (DDZ) and Leibniz Institute for Diabetes Research, Düsseldorf, NRW, DE

^30^ Wellcome Sanger Institute, Genome Campus, Hinxton, CB10 1RQ, UK

^31^ Department of Genetics, University of Cambridge, Cambridge, CB2 3EH, UK

^32^ Institute for Medical Biometry and Bioinformatics, Medical Faculty and University Hospital Düsseldorf, Heinrich Heine University, Düsseldorf, NRW, DE

^33^ Howard Hughes Medical Institute, Chevy Chase, MD 20815, USA

^34^ ISEM, Univ Montpellier, CNRS, IRD, Montpellier, FR

^35^ Institut Universitaire de France, Paris, FR

^36^ Department of Computer Science and Engineering, University of California San Diego, La Jolla, CA 92093, USA

^37^ Department of Bioethics & Humanities, University of Washington School of Medicine, Seattle, WA 98195, USA

^38^ Department of Genetics, Epigenetics Institute, Perelman School of Medicine, University of Pennsylvania, Philadelphia, PA 19104, USA

^39^ Department of Data Science, Dana-Farber Cancer Institute, Boston, MA 02215, USA

^40^ Department of Anthropology, University of Kansas, Lawrence, KS 66045, USA

^41^ School of Health Sciences, University of Manchester, Manchester M13 9PL, UK

^42^ Traditional, ancestral and unceded territory of the Gabrielino/Tongva peoples, Institute for Society & Genetics, University of California, Los Angeles, Los Angeles, CA 90095, USA

^43^ Traditional, ancestral and unceded territory of the Gabrielino/Tongva peoples, Institute for Precision Health, David Geffen School of Medicine, University of California, Los Angeles, Los Angeles, CA 90095, USA

^44^ Traditional, ancestral and unceded territory of the Gabrielino/Tongva peoples, Division of General Internal Medicine & Health Services Research, David Geffen School of Medicine, University of California, Los Angeles, Los Angeles, CA 90095, USA

^45^ Department of Electrical Engineering and Computer Science, University of California, Berkeley, Berkeley, CA 94720, USA

^46^ Department of Biomedical Data Science, Stanford University School of Medicine, Stanford, CA 94305, USA

^47^ Medical and Population Genomics Lab, Sidra Medicine, Doha, Qatar

^48^ Department of Molecular Cell and Developmental Biology, University of California, Santa Cruz, CA, USA

^49^ Montreal Heart Institute, Montréal, QC, Canada

^50^ Department of Pediatrics, University of California San Diego, La Jolla, CA 92093, USA

^51^ Center for Genomic Health, Yale University School of Medicine, New Haven, CT 06510, USA

^52^ Department of Genetics, Yale University School of Medicine, New Haven, CT 06510, USA

^53^ Department of Biomedical Informatics, Harvard Medical School, Boston, MA 02115, USA

^54^ Department of Evolution and Ecology and the Center for Population Biology, University of California, One Shields, Davis, CA 95616, USA

^55^ Department of Computer Science, Johns Hopkins University, Baltimore, MD 21218, USA

^56^ Department of Pediatrics, Division of Genetics, School of Medicine, University of California, Irvine, CA 92697, USA

^57^ Sun Yat-sen University, Guangzhou, China

^58^ Edison Family Center for Genome Sciences & Systems Biology, Washington University School of Medicine, St. Louis, MO 63110, USA

^59^ Department of Biology and Center for Medical Genomics, Penn State University, University Park, PA 16802, USA

^60^ Division of Medical Genetics, Department of Medicine, University of Washington School of Medicine, Seattle, WA 98195, USA

^61^ The Jackson Laboratory for Genomic Medicine, Farmington, CT 06032, USA

^62^ Department of Biology, Penn State University, University Park, PA 16802, USA

^63^ Department of Biomedical Science, College of Health Sciences, Qatar University, Doha, Qatar

^64^ Department of Genetic Medicine, Weill Cornell Medicine-Qatar, Doha, Qatar

^65^ IRSD - Digestive Health Research Institute, University of Toulouse, INSERM, INRAE, ENVT,

UPS, Toulouse, FR

^66^ MATCH biosystems, S.L., Elche, Spain

^67^ Universidad Miguel Hernández de Elche, Elche, Spain

^68^ Department of Computational Biology and Medical Sciences, The University of Tokyo, Kashiwa, Chiba 277-8561, Japan

^69^ Department of Computer Science, University of Pisa, Pisa, Italy

^70^ Law School, University of Wisconsin-Madison, Madison, WI 53706, USA

^71^ Morgridge Institute for Research, Madison, WI 53715, USA

^72^ Institute of Genetics and Biomedical Research, UoS of Milan, National Research Council, Milan, Italy

^73^ Department of Biomedical Engineering, Johns Hopkins University, Baltimore, MD 21218, USA

^74^ Department of Genetic Medicine, Johns Hopkins University School of Medicine, Baltimore, MD 21205, USA

^75^ Genome Biology Unit, European Molecular Biology Laboratory (EMBL), Heidelberg, DE

^76^ Institute for Molecular Medicine Finland, Helsinki Institute of Life Science, University of Helsinki, Helsinki, Finland

^77^ The Center for Bio- and Medical Technologies, Moscow, RUS

^78^ Centre for Biomedical Research and Technology, HSE University, Moscow, RUS

^79^ Department of Biology, Johns Hopkins University, Baltimore, MD 21218, USA

^80^ Coriell Institute for Medical Research, Camden, NJ 08103, USA

^81^ University of Amsterdam, Amsterdam, Netherlands

^82^ School of Clinical Medicine, University of Cambridge, Cambridge, CB2 0SP, UK

^83^ Center for Genomic Discovery, Mohammed Bin Rashid University, Dubai Health, UAE

^84^ Dubai Health Genomic Medicine Center, Dubai Health, UAE

^85^ GenomeArc Inc, Mississauga, ON, Canada

^86^ Department of Biology and Biotechnologies “Charles Darwin”, University of Rome “La Sapienza”, Rome 00185, IT

^87^ Department of Human Genetics and Utah Center for Genetic Discovery, University of Utah, Salt Lake City, UT, USA

^88^ Center for Genomics, Loma Linda University School of Medicine, Loma Linda, CA 92350, USA

^89^ PacBio, Menlo Park, CA 94025, USA

^90^ The first affiliated hospital of Xi’an Jiaotong University, Xi’an Jiaotong University, Xi’an, Shaanxi, 710049, China

## Human Genome Structural Variation Consortium Authors

Hufsah Ashraf^1,2^, Peter A. Audano^3^, Marcelo Ayllon^4^, Andrey Azov^5^, Parithi Balachandran^3^, Anna O. Basile^6^, Christine R. Beck^3,7^, Marc Jan Bonder^8,9,10^, Lucy Brooks^5^, Marta Byrska-Bishop^6^, Mark J.P. Chaisson^11^, Zechen Chong^12^, Wayne E. Clarke^6,13^, André Corvelo^6^, Jonathan Crabtree^14^, Scott E. Devine^14^, Peter Ebert^15,2^, Jana Ebler^1,2^, Evan E. Eichler^4,16^, Aine Fairbrother-Browne^5^, Chia-Hsuan Fan^12^, Mallory Freeberg^5^, Shenghan Gao^17^, Mark B. Gerstein^18,19^, Bida Gu^11^, Pille Hallast^3^, Patrick Hasenfeld^20^, Mir Henglin^1,2^, Kendra Hoekzema^4^, Kaili Hu Hu^12^, Sarah Hunt^5^, Matthew Jensen^18,19^, Yunzhe Jiang^18,19^, Tomoya Kanno^21,22^, Kwondo Kim^7^, Miriam K. Konkel^23,24^, Jan O. Korbel^20^, Youngjun Kwon^4^, Peter M. Lansdorp^25,26^, Charles Lee^7^, Tiffany Leung^25^, Jiaqi Li^18,19^, Jiadong Lin^4^, Mark Loftus^7^, Glennis A. Logsdon^17^, Tobias Marschall^1,2^, Gianni V. Martino^23^, Ryan E. Mills^27^, Yulia Mostovoy^28^, Katherine M. Munson^4^, Giuseppe Narzisi^6^, Lingbin Ni^4^, Keisuke K. Oshima^17^, Carolyn Paisie^3^, Samarendra Pani^1,2^, Zishan Peng^12^, David Porubsky^4^, Timofey Prodanov^1,2^, Keon Rabbani^11^, Valmik Ranparia^11^, Tobias Rausch^20^, Luyao Ren^4,16^, Xinghua Shi^21,22^, Yuwei Song^12^, Kaitlyn Sun^4^, Likhitha Surapaneni^5^, Michael E. Talkowski^29,30,31^, Vasiliki Tsapalou^20^, Andres Veidenberg^5^, Feyza Yilmaz^7^, DongAhn Yoo^4^, Xuefang Zhao^29,30,31^, Weichen Zhou^27^, Michael C. Zody^6^

### Affiliations

^1^ Institute for Medical Biometry and Bioinformatics, Medical Faculty and University Hospital Düsseldorf, Heinrich Heine University, Düsseldorf, Germany

^2^ Center for Digital Medicine, Heinrich Heine University, Düsseldorf, Germany

^3^ The Jackson Laboratory for Genomic Medicine, Farmington, CT, USA

^4^ University of Washington School of Medicine, Department of Genome Sciences, Seattle, WA, USA

^5^ European Molecular Biology Laboratory, Wellcome Genome Campus, European Bioinformatics Institute, Cambridge, UK

^6^ New York Genome Center, New York, NY, USA

^7^ The University of Connecticut Health Center, Farmington, CT, USA

^8^ Department of Genetics, University of Groningen, University Medical Center Groningen, Groningen, The Netherlands

^9^ Oncode Institute, Utrecht, The Netherlands

^10^ Division of Computational Genomics and Systems Genetics, German Cancer Research Center, Heidelberg, Germany

^11^ Department of Quantitative and Computational Biology, University of Southern California, Los Angeles, CA, USA

^12^ Department of Biomedical Informatics and Data Science, Heersink School of Medicine, University of Alabama, Birmingham, AL, USA

^13^ Outlier Informatics Inc., Saskatoon, SK S7H 1L4, Canada

^14^ Institute for Genome Sciences, University of Maryland School of Medicine, Baltimore, MD 21201 USA

^15^ Core Unit Bioinformatics, Medical Faculty and University Hospital Düsseldorf, Heinrich Heine University, Düsseldorf, Germany

^16^ Howard Hughes Medical Institute, University of Washington, Seattle, WA, USA

^17^ Department of Genetics, Epigenetics Institute, Perelman School of Medicine, University of Pennsylvania, Philadelphia, PA, USA

^18^ Department of Molecular Biophysics and Biochemistry, Yale University, New Haven, CT, USA

^19^ Program in Computational Biology and Bioinformatics, Yale University, New Haven, CT, USA

^20^ European Molecular Biology Laboratory (EMBL), Genome Biology Unit, Heidelberg, Germany

^21^ Department of Computer and Information Sciences, College of Science and Technology, Temple University, Philadelphia, PA, USA

^22^ Institute for Genomics and Evolutionary Medicine, Temple University, Philadelphia, PA, USA

^23^ Clemson University, Department of Genetics & Biochemistry, Clemson, SC, USA

^24^ Center for Human Genetics, Clemson University, Greenwood, SC, USA

^25^ Terry Fox Laboratory, BC Cancer Agency, Vancouver, BC, V5Z 1L3, Canada

^26^ Department of Medical Genetics, University of British Columbia, Vancouver, BC, V6T 1Z4, Canada

^27^ Department of Computational Medicine & Bioinformatics, University of Michigan, MI, 48109, USA

^28^ Cardiovascular Research Institute and Institute for Human Genetics, UCSF School of Medicine, CA, USA

^29^ Program in Medical and Population Genetics and Stanley Center for Psychiatric Research, Broad Institute of MIT and Harvard, Cambridge, MA, USA

^30^ Center for Genomic Medicine, Massachusetts General Hospital, Boston, MA, USA

^31^ Department of Neurology, Massachusetts General Hospital and Harvard Medical School, Boston, MA, USA

