## Extended Figures for "Pangenome discovery and characterization of human protein-coding duplicated genes"

### EXTENDED DATA FIGURES

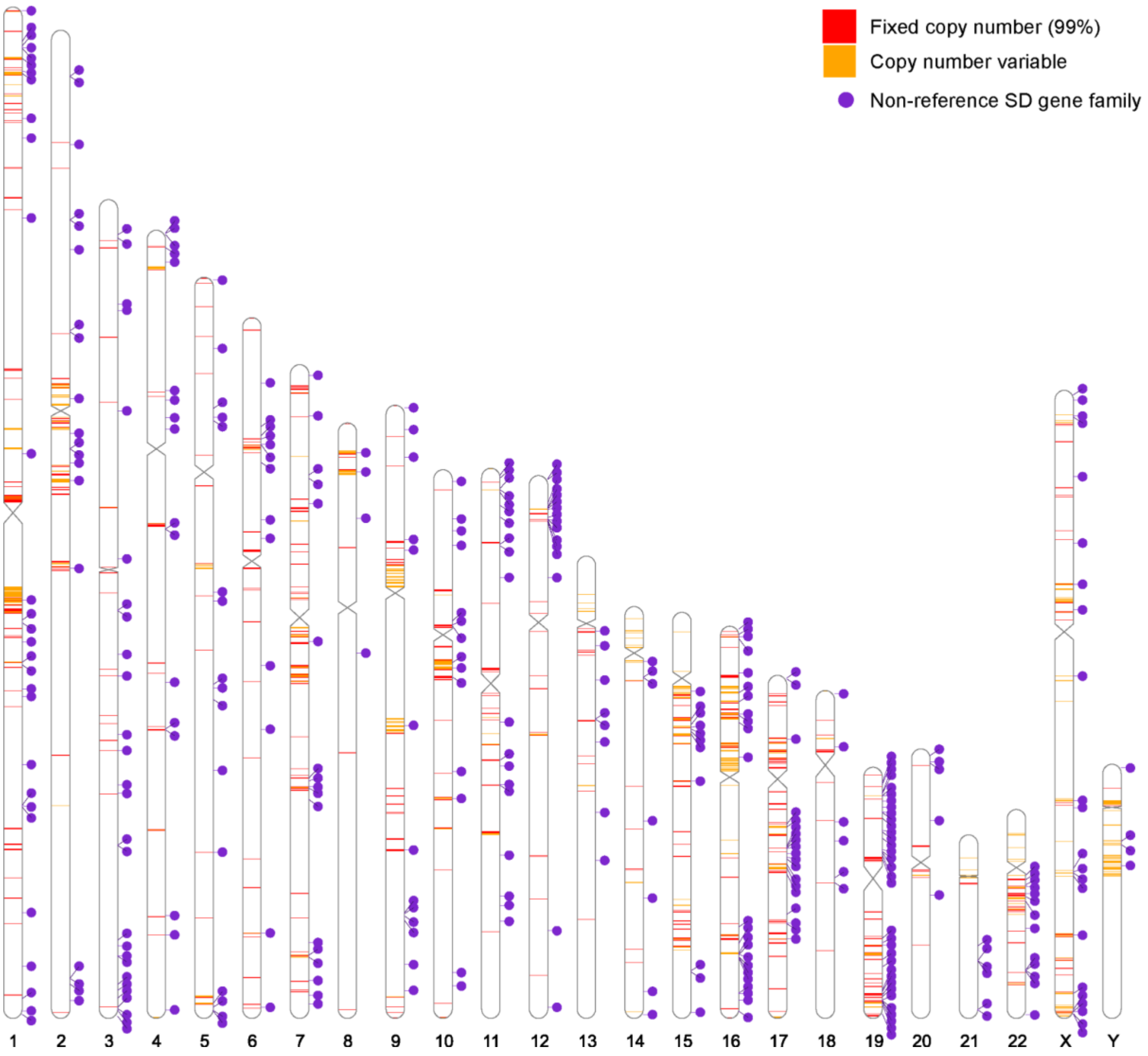


**Extended Data Fig. 1 | Chromosomal distribution of SD gene copy number categories in T2T-CHM13.** Colored bands show the positions of SD genes along each chromosome. Red denotes fixed SD genes, defined as genes present in more than 99% of haplotypes at a stable copy number of one, whereas orange denotes copy number variable SD genes. Purple circles, connected by lines to their corresponding loci, indicate non-reference SD gene families—paralog groups represented by a single, nonduplicated locus in the T2T-CHM13 reference but duplicated in one or more other haplotypes across the pangenome.

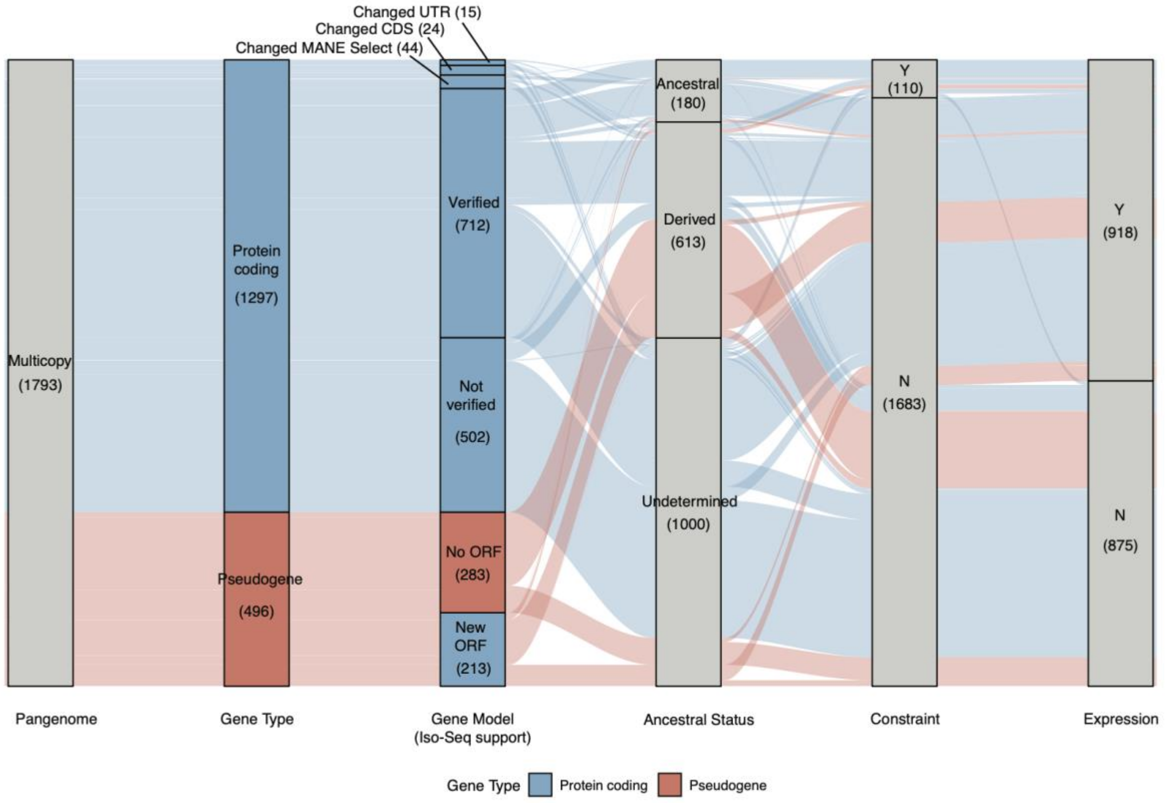


**Extended Data Fig. 2 | Overview of 1793 multi-copy SD gene characterization/reclassification.** The alluvial plot shows the refinement of 1793 SD genes that are absent from the reference genome but are too similar to be assigned to distinct phylogenetic clades. The legend is the same as in Fig. 1C.

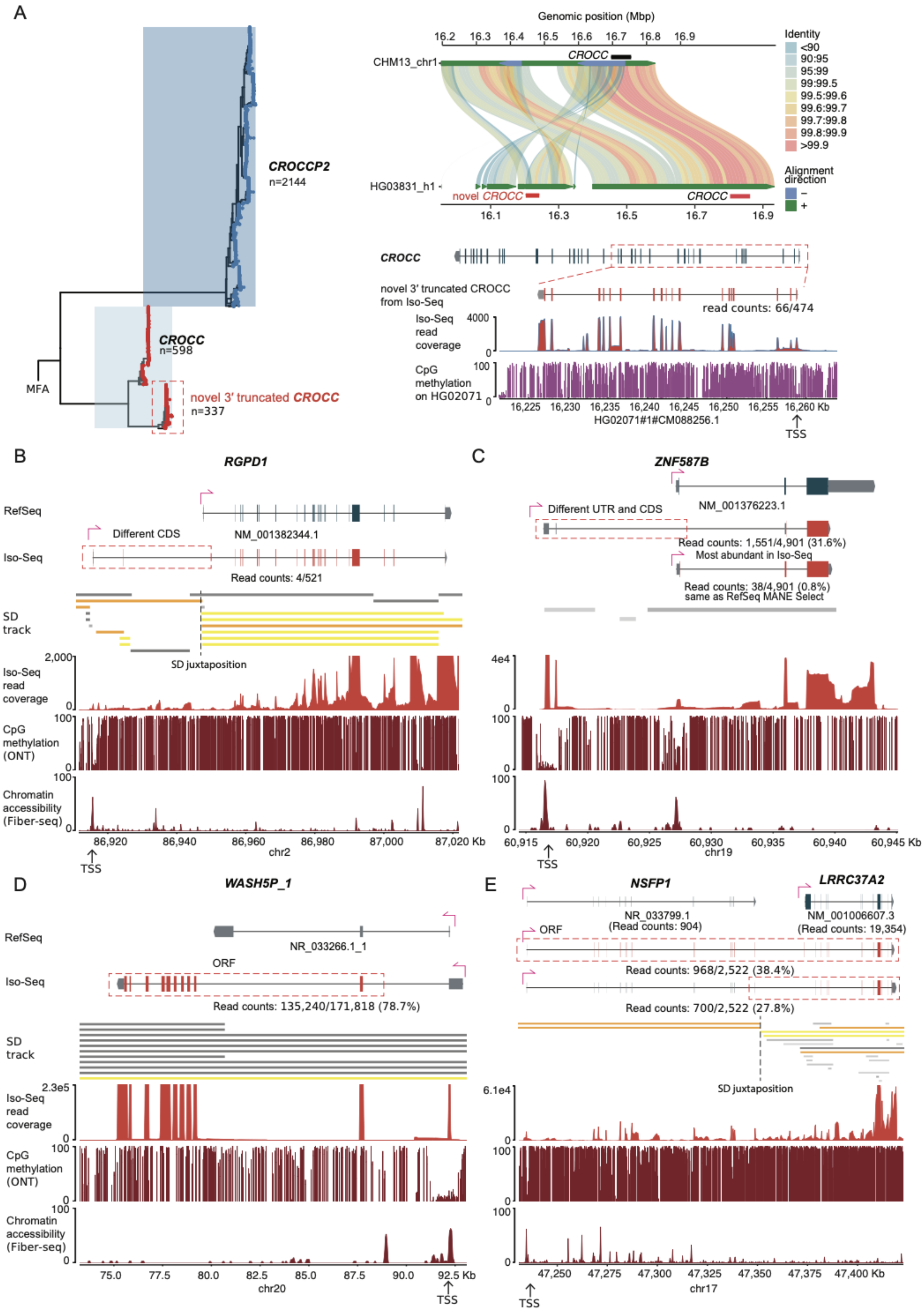
A
CROCCP2
n=2144
CROCC
n=598
novel 3' truncated CROCC
n=337
MFA
Genomic position (Mbp)
16.2 16.3 16.4 16.5 16.6 16.7 16.8 16.9
CHM13_chr1
CROCC
HG03831_h1
novel CROCC
CROCC
16.1 16.3 16.5 16.7 16.9
Identity
<90
90:95
95:99
99:99.5
99.5:99.6
99.6:99.7
99.7:99.8
99.8:99.9
>99.9
Alignment direction
–
+
CROCC
novel 3' truncated CROCC from Iso-Seq
read counts: 66/474
Iso-Seq read coverage
4000
0
CpG methylation on HG02071
100
0
16,225 16,230 16,235 16,240 16,245 16,250 16,255 16,260 Kb
HG02071#1#CM088256.1
TSS
B
RGPD1
RefSeq
Different CDS
NM_001382344.1
Iso-Seq
Read counts: 4/521
SD track
SD juxtaposition
2,000
Iso-Seq read coverage
0
100
CpG methylation (ONT)
0
100
Chromatin accessibility (Fiber-seq)
0
86,920 86,940 86,960 86,980 87,000 87,020 Kb
chr2
TSS
C
ZNF587B
Different UTR and CDS
NM_001376223.1
Read counts: 1,551/4,901 (31.6%)
Most abundant in Iso-Seq
Read counts: 38/4,901 (0.8%)
same as RefSeq MANE Select
4e4
0
100
0
100
0
60,915 60,920 60,925 60,930 60,935 60,940 60,945 Kb
chr19
TSS
D
WASH5P_1
RefSeq
NR_033266.1_1
ORF
Iso-Seq
Read counts: 135,240/171,818 (78.7%)
SD track
2.3e5
Iso-Seq read coverage
0
100
CpG methylation (ONT)
0
100
Chromatin accessibility (Fiber-seq)
0
75.0 77.5 80.0 82.5 85.0 87.5 90.0 92.5 Kb
chr20
TSS
E
NSFP1
LRRC37A2
NR_033799.1
(Read counts: 904)
NM_001006607.3
(Read counts: 19,354)
ORF
Read counts: 968/2,522 (38.4%)
Read counts: 700/2,522 (27.8%)
SD juxtaposition
6.1e4
0
100
0
100
0
47,250 47,275 47,300 47,325 47,350 47,375 47,400 Kb
chr17
TSS

**Extended Data Fig. 3 | Validation of RefSeq gene annotations using Iso-Seq.**
**(A)** New CROCC-derived gene models were identified from clades that clustered without T2T-CHM13 reference genes. The left panel shows a phylogenetic tree rooted with MFA as the outgroup. The upper-right panel shows an SVbyEye plot identifying an additional truncated CROCC copy in the HG03831_h1 assembly relative to T2T-CHM13. The lower-right panel shows gene models identified by mapping Iso-Seq reads to the HG02071_h1 assembly, with Iso-Seq read coverage and CpG methylation tracks shown below. **(B, C)** Selected examples of gene models identified by Iso-Seq that differ from RefSeq annotations, including alternative 5′ promoters: *RGPD1* in **(B)** and *ZNF587B* in **(C)**. **(D)** Selected example, *WASH5P_1*, of the reclassification of a pseudogene as a protein-coding gene. **(E)** Selected example of a gene fusion involving *NSFP1* and *LRRC37A2*. The four tracks below show SD track, Iso-Seq read coverage, CpG methylation in T2T-CHM13, and chromatin actuation in T2T-CHM13, indicating promoter activity.

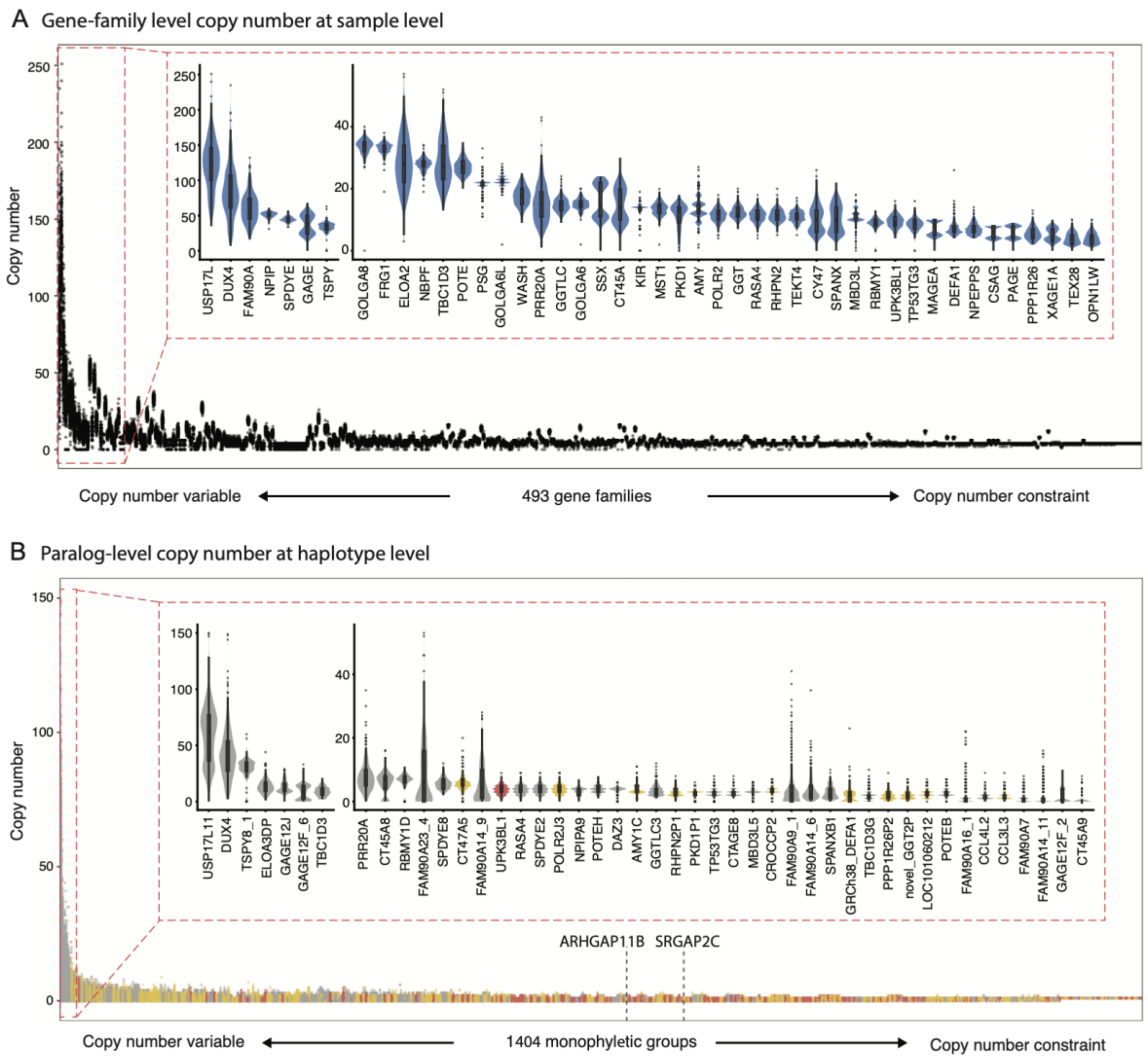


**Extended Data Fig. 4 | Gene-family-level and haplotype-level SD gene copy number. (A)** Gene-family-level (n = 493) copy number at sample level, ranked by copy number standard deviation. The red box indicates the 45 most copy number variable genes. Gene copies located in assembly error regions were excluded. **(B)** Paralog-level (n = 1,404) copy number across haplotypes, ranked by copy number standard deviation. Red color represents ancestral genes, yellow color represents derived genes, and gray color represents genes with undetermined ancestral status. The red box indicates the 45 most copy number variable genes.
