## Supplementary Figures for "Pangenome discovery and characterization of human protein-coding duplicated genes"

### Supplementary Information

### Glossary

**Segmental duplication (SD) genes**: Genes with >30% of the gene body/transcript located in an SD pairwise alignment (>90% & >1 kbp). These can be further subdivided into **ancestral** (progenitor copy typically syntenic in location in other nonhuman primates where SD has not occurred) or **derived** (duplicate copies that are more likely to be lineage-specific). A subset of SD genes cannot be assigned to an ancestral or derived status and are classified as **undetermined**.

**Paralog**: A homologous (gene) copy that arose by duplication in a genome.

**Paragroup**: Multiple gene copies are grouped within a broader clade but cannot be assigned to a distinct phylogenetic group because they are too similar either as a result of recent duplication or interlocus gene conversion.

**Reference SD genes**: Genes annotated as mapping to SDs in the human reference genome, T2T-CHM13. These may be copy number variable with fixed copy number in the human pangenome.

**Non-reference SDs**: Gene that maps to an SD gene family in T2T-CHM13 but is phylogenetically distinct from any copy in the reference.

**Increased copy number**: Additional duplicate genes in a human haplotype when compared to a reference but which cannot be phylogenetically distinguished.

**Non-reference SD gene family**: Gene that maps to a unique sequence in the T2T-CHM13 reference genome but duplicated in a subset of human genomes and by definition is copy number variable. These are further classified as singleton if the duplication is only observed in one human genome, doubleton if observed twice, and polymorphic if observed two or more times.

### Supplementary Figures

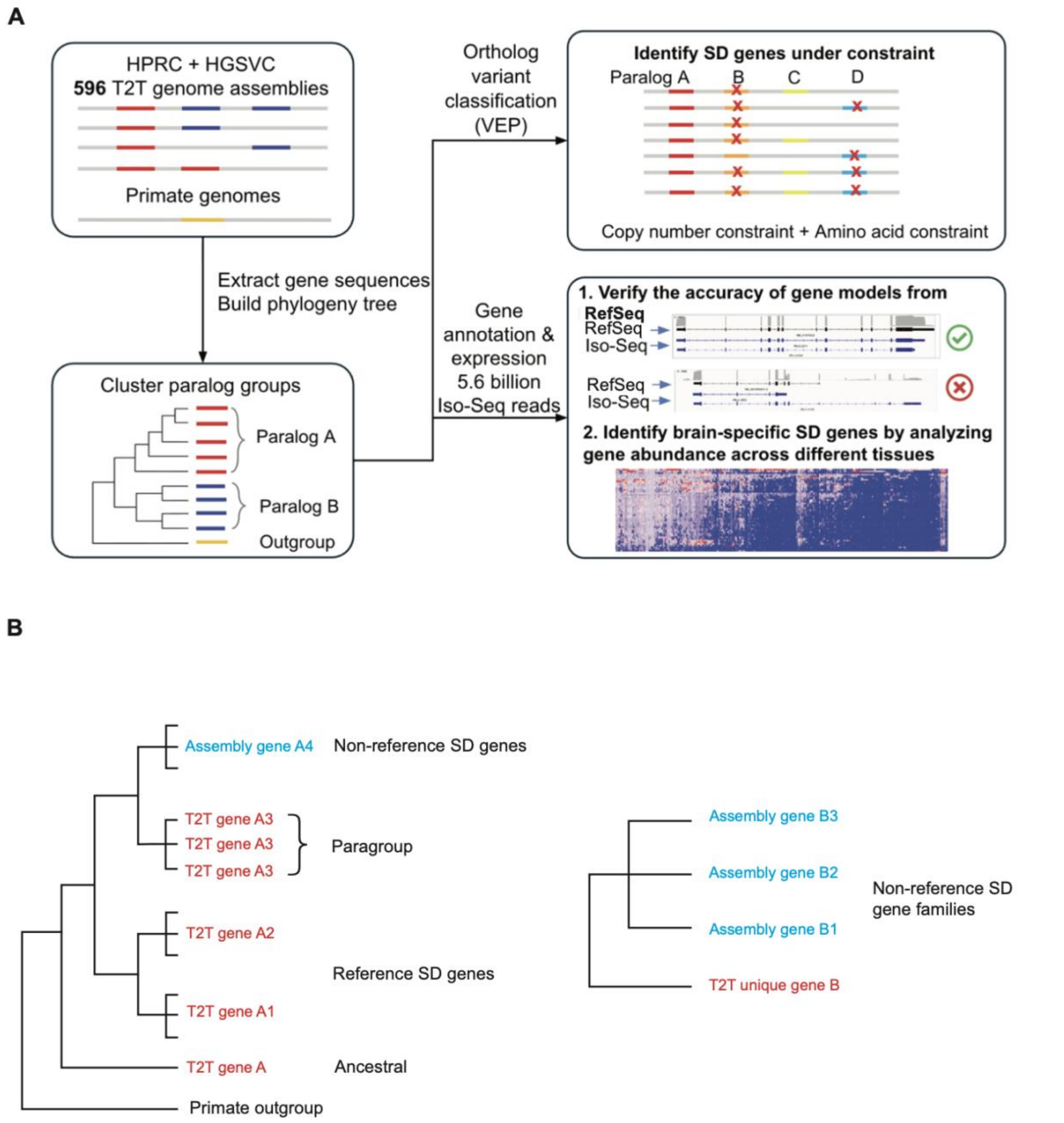


**Supplementary Fig. 1 | Study workflow and glossary. (A)** Overview of the research strategy and analytical pipeline. Abbreviations: HPRC, Human Pangenome Reference Consortium; HGSVC, Human Genome Structural Variation Consortium; T2T, Telomere-to-Telomere; VEP, Ensembl Variant Effect Predictor. **(B)** Conceptual diagram illustrating the phylogenetic terms used for SD gene classification.

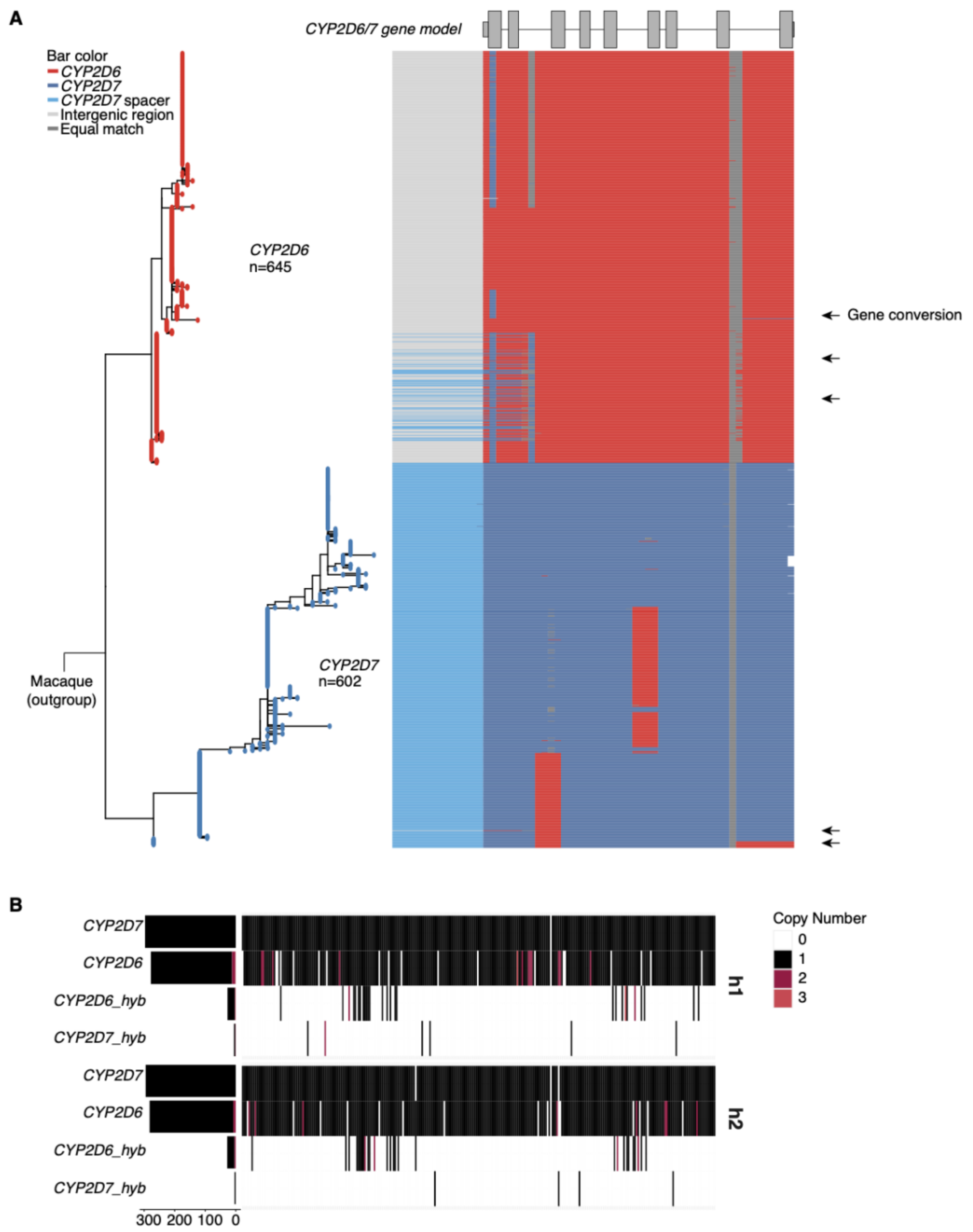


**Supplementary Fig. 2 | Gene conversion in *CYP2D6* and *CYP2D7*.** Potential interlocus gene conversion (IGC) events within a gene family can often be inferred from phylogenetic tree topology. For example, a staircase-like pattern near the root of a clade or unusually long branches within a clade may indicate gene conversion, superimposing these patterns onto the topology of each human SD gene family. **(A)** Phylogenetic tree of *CYP2D6* and *CYP2D7* with gene conversion events. The heatmap shows the best-matching regions within genes relative to the T2T-CHM13 reference genome, with arrows indicating inferred gene conversion tracts. **(B)** Copy number of *CYP2D6* and *CYP2D7* across both haplotypes for each HPRC and HGSVC individual.

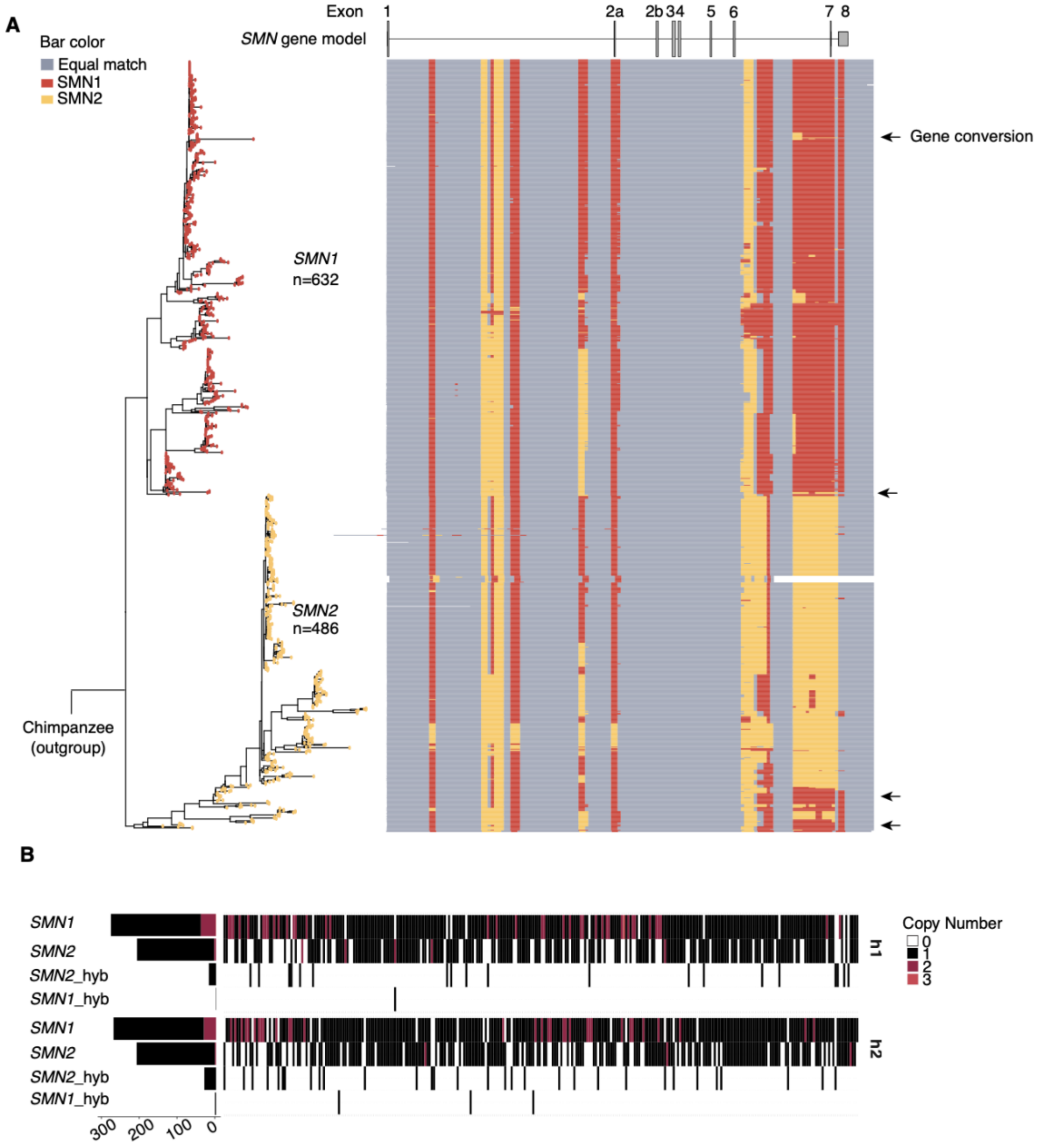


**Supplementary Fig. 3 | Gene conversion in *SMN1* and *SMN2*.** Same legend as Supplementary Fig. 2.

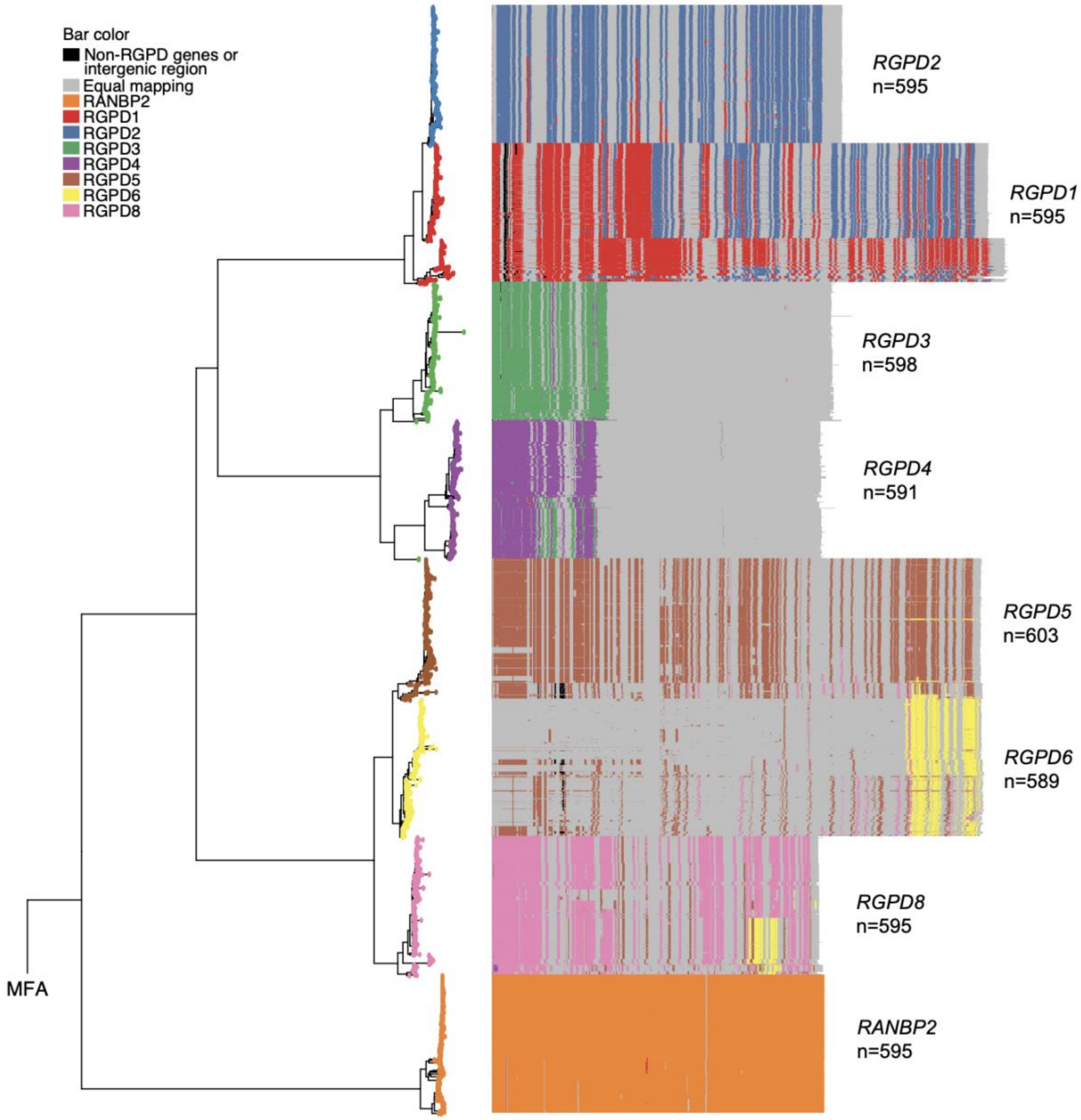


**Supplementary Fig. 4 | Gene conversion events identified in the *RGPD* gene family.** Same legend as Supplementary Fig. 2.

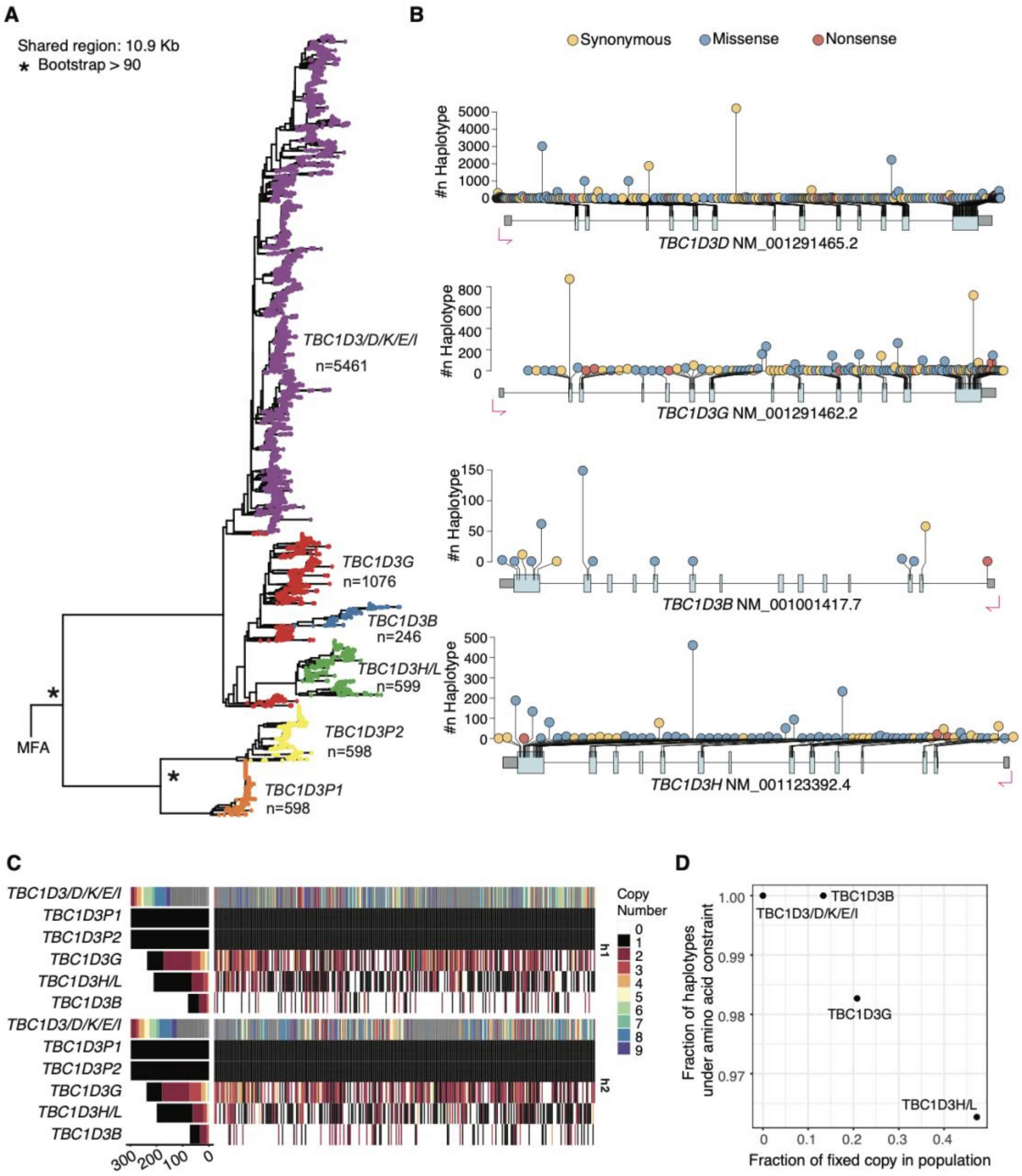


**Supplementary Fig. 5 | Paralog grouping, copy number, and variants for core duplications in the *TBC1D3* gene family. (A)** Population-based phylogenetic tree rooted using macaque (MFA) as the outgroup. Paralog grouping based on a 1.5× allelic-variation threshold. Integer values represent copy numbers in 596 HPRC and HGSVC human genomes, T2T-CHM13, and GRCh38. **(B)** Amino acid constraint showing synonymous (yellow), amino acid substitutions (blue) and stop codon (red) polymorphisms predicted using VEP based on a gene model with long-read transcriptome support. **(C)** Heatmap summarizes the copy number of each paralog across both haplotypes for each individual haplotype, with gene deletions (white) and duplications (red) indicated. **(D)** Scatter plot of the constraint map for *TBC1D3* gene family.

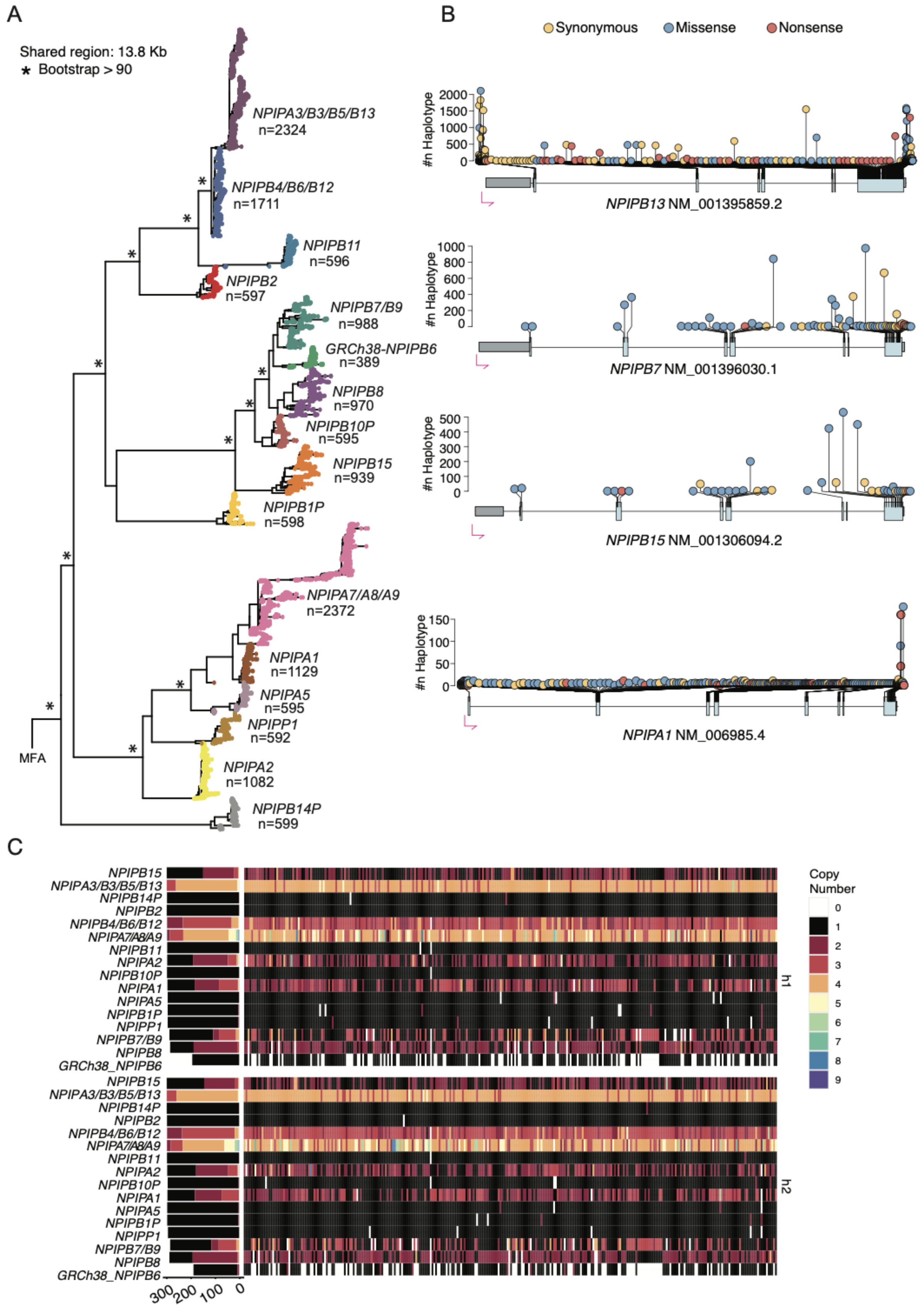


**Supplementary Fig. 6 | Paralog grouping, copy number, and variants for core duplications in the *NPIP* gene family.** Same legend as Supplementary Fig. 5.

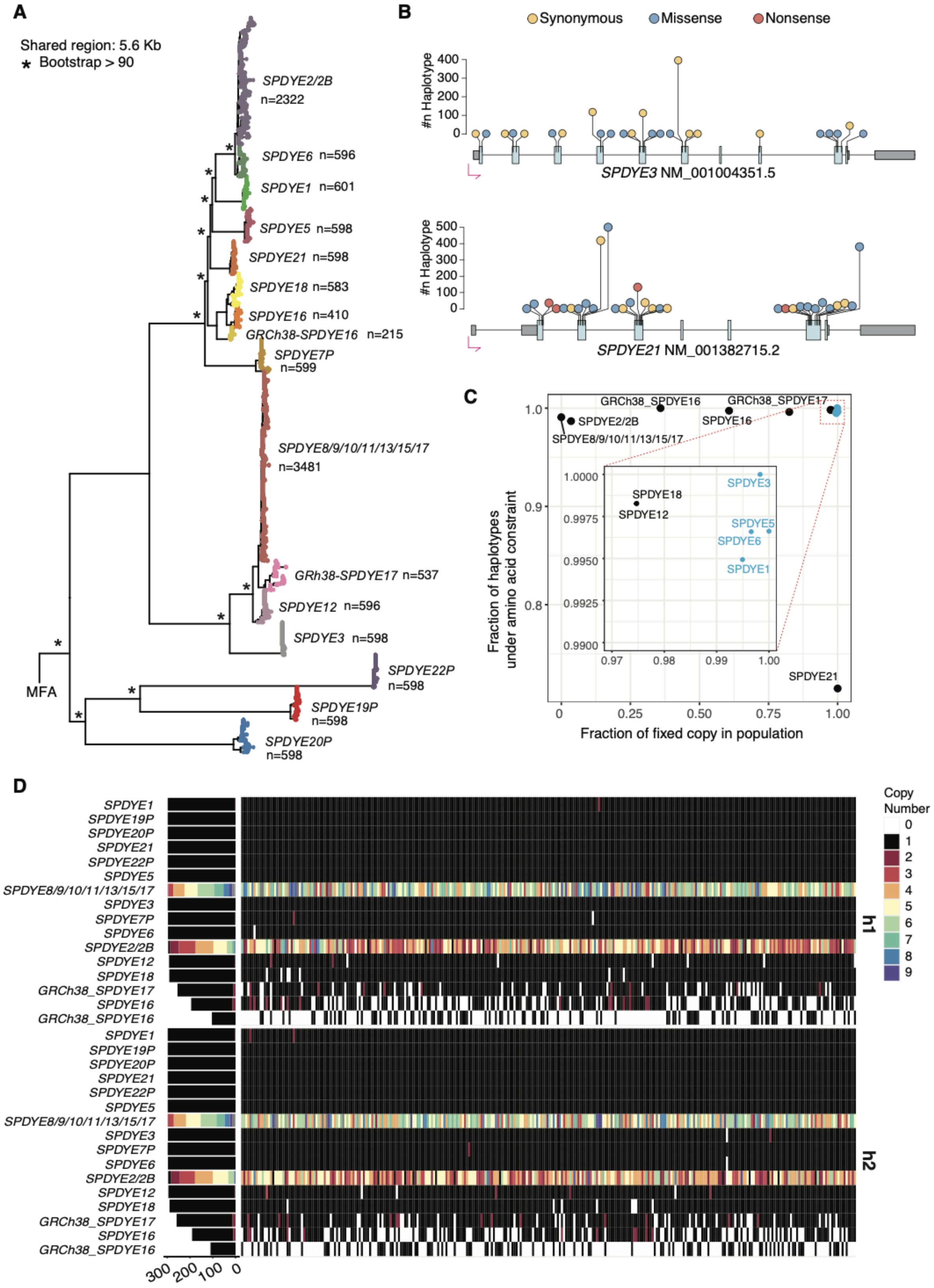


**Supplementary Fig. 7 | Paralog grouping, copy number, and variants for core duplications in the *SPDYE* gene family.** Same legend as Supplementary Fig. 5.

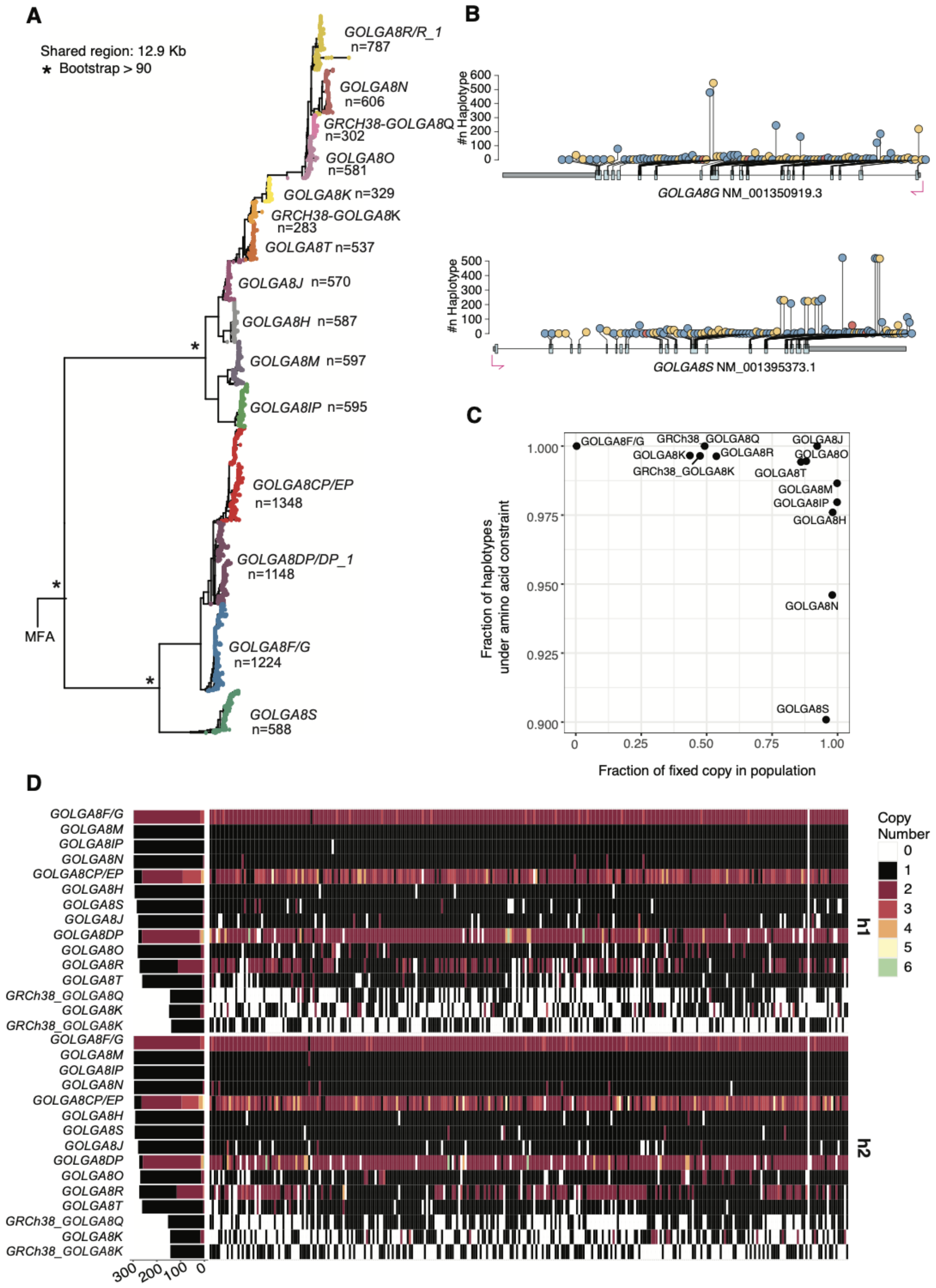


**Supplementary Fig. 8 | Paralog grouping, copy number, and variants for core duplications in the *GOLGA8* gene family.** Same legend as Supplementary Fig. 5.

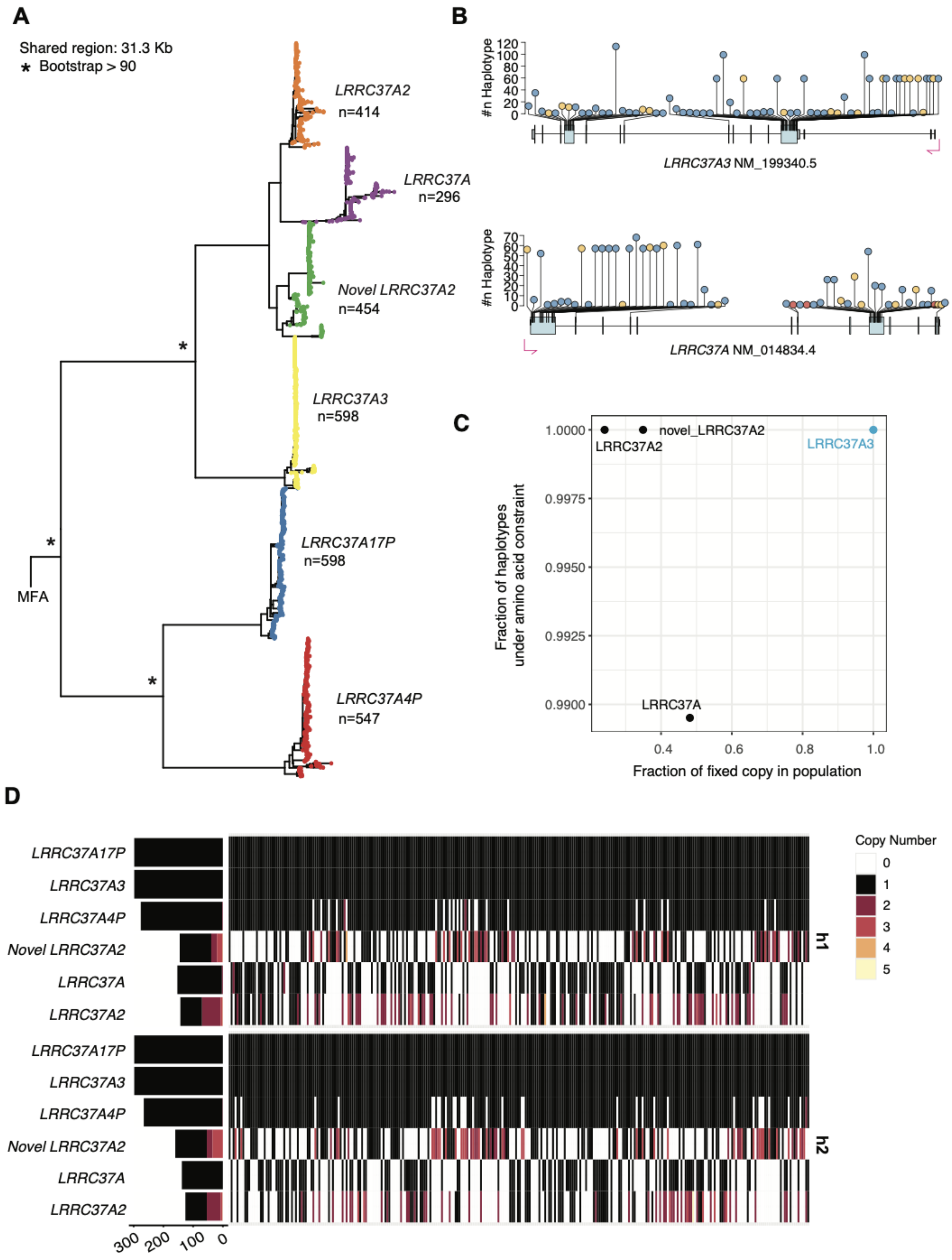


**Supplementary Fig. 9 | Paralog grouping, copy number, and variants for core duplications in the *LRRC37A* gene family.** Same legend as Supplementary Fig. 5.

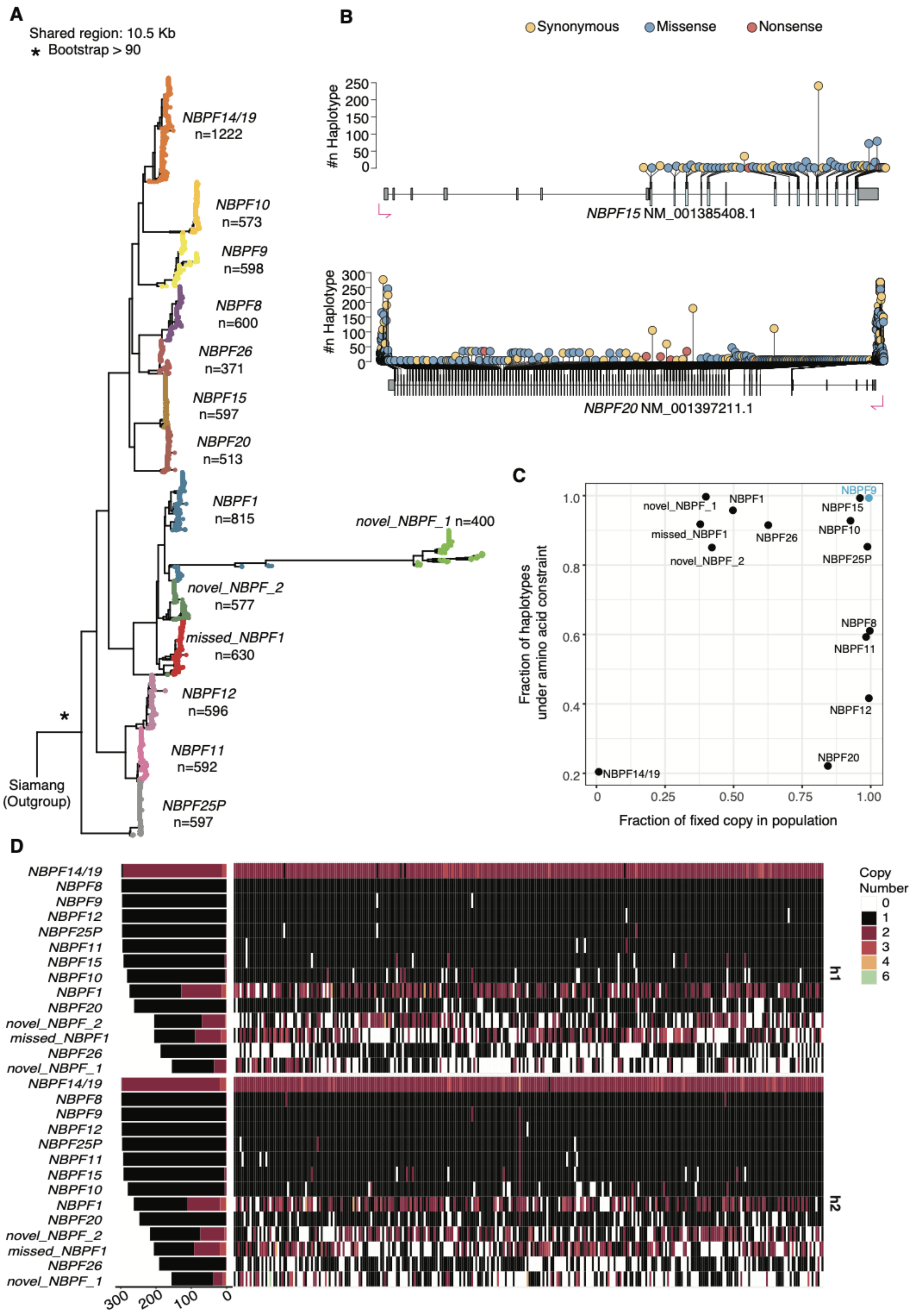


**Supplementary Fig. 10. Paralog grouping, copy number, and variants for core duplications in the *NBPF* gene family.** Same legend as Supplementary Fig. 5.

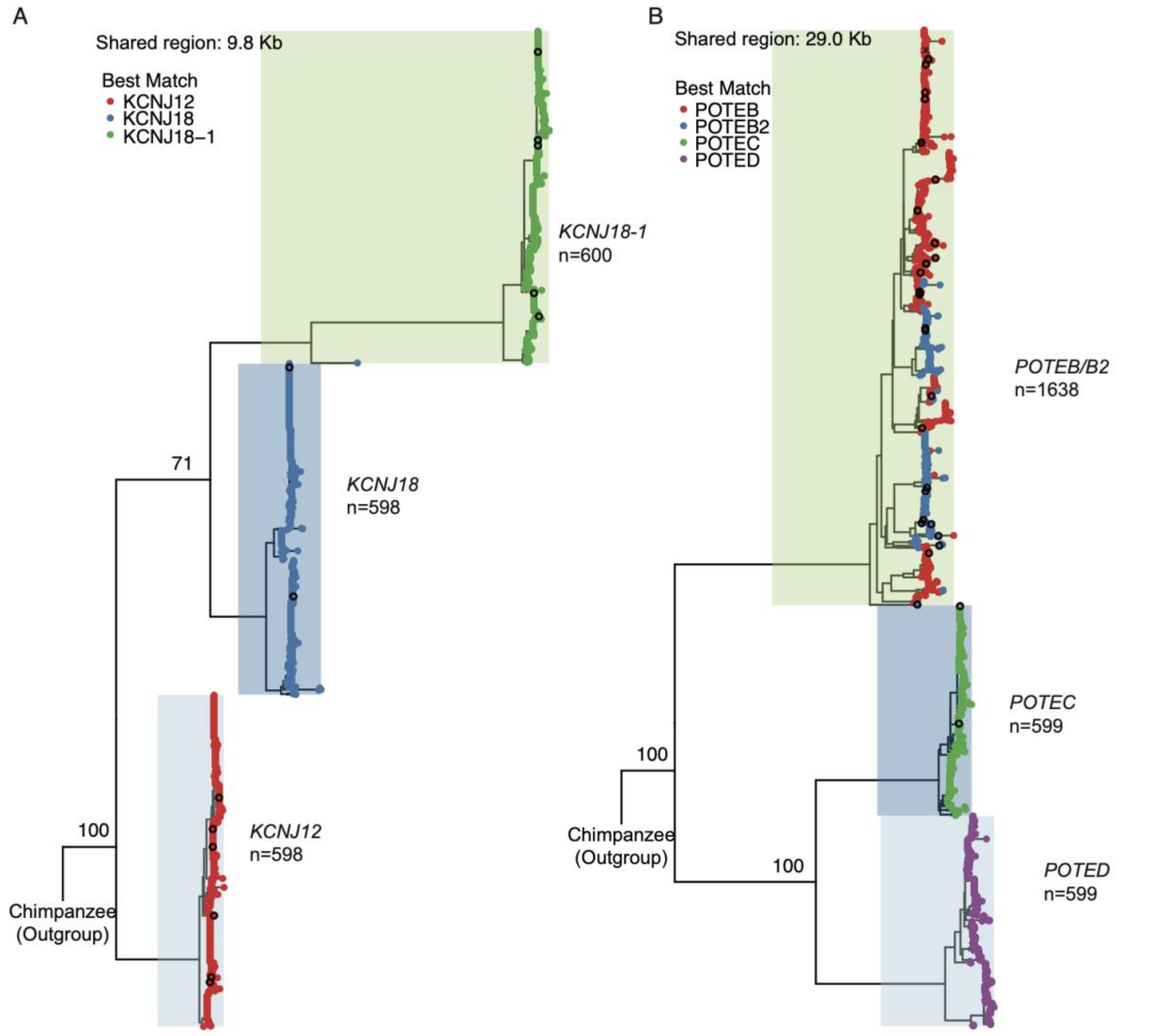


**Supplementary Fig. 11 | Tree topology to curate gene annotation. (A)** *KCNJ18-1* is annotated as a duplicated copy of *KCNJ18* but falls into a different clade. **(B)** *POTEB* (ENSG00000233917) and *POTEB2* (ENSG00000230031) paralogs cluster in a single clade. The dot color indicates the best match identified by minimap2.

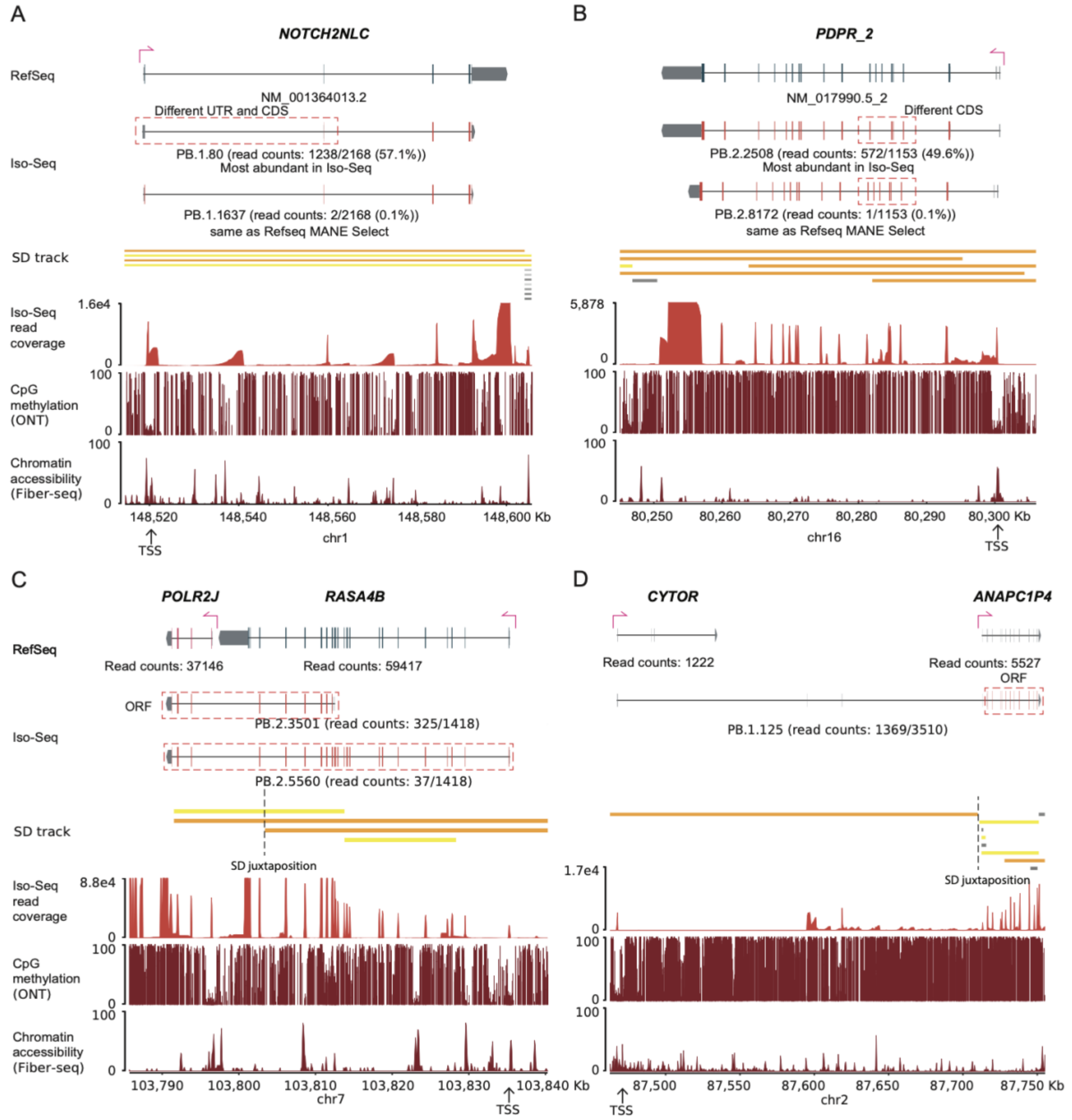


**Supplementary Fig. 12 | Gene models identified from Iso-Seq.** Iso-Seq–based corrections to MANE Select RefSeq annotations for **(A)** *NOTCH2NLC* and **(B)** *PDPR_2*. Iso-Seq–identified gene fusions arising from the juxtaposition of two SDs: **(C)** *POLR2J–RASA4B* and **(D)** *CYTOR–ANAPC1P4*.

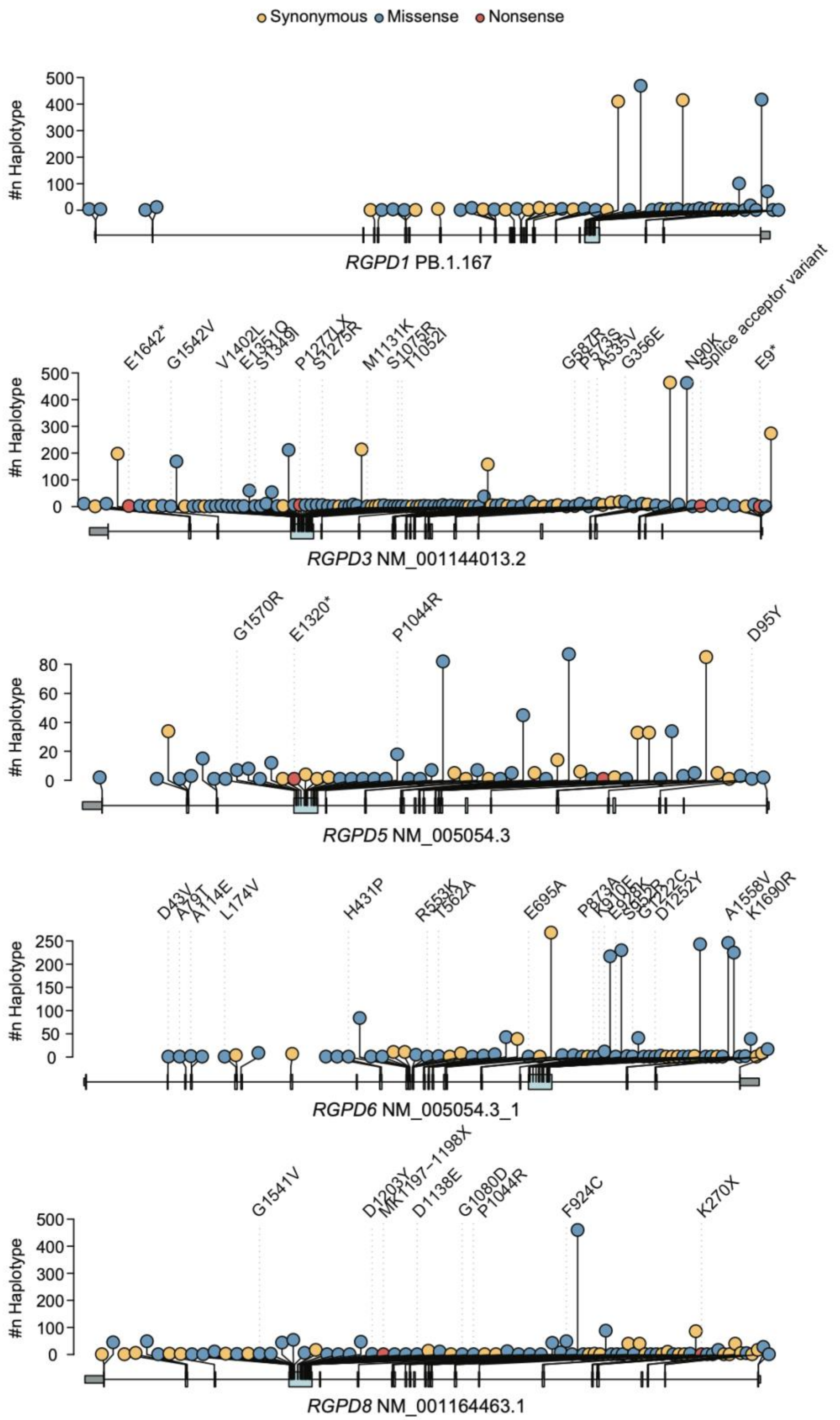


**Supplementary Fig. 13 | Variants identified from *RGPD1*, *RGPD3*, *RGPD5*, *RGPD6*, and *RGPD8*.** Amino acid constraint showing synonymous (yellow), amino acid substitutions (blue), and stop codon (red) polymorphisms predicted using VEP based on a gene model with long-read transcriptome support.

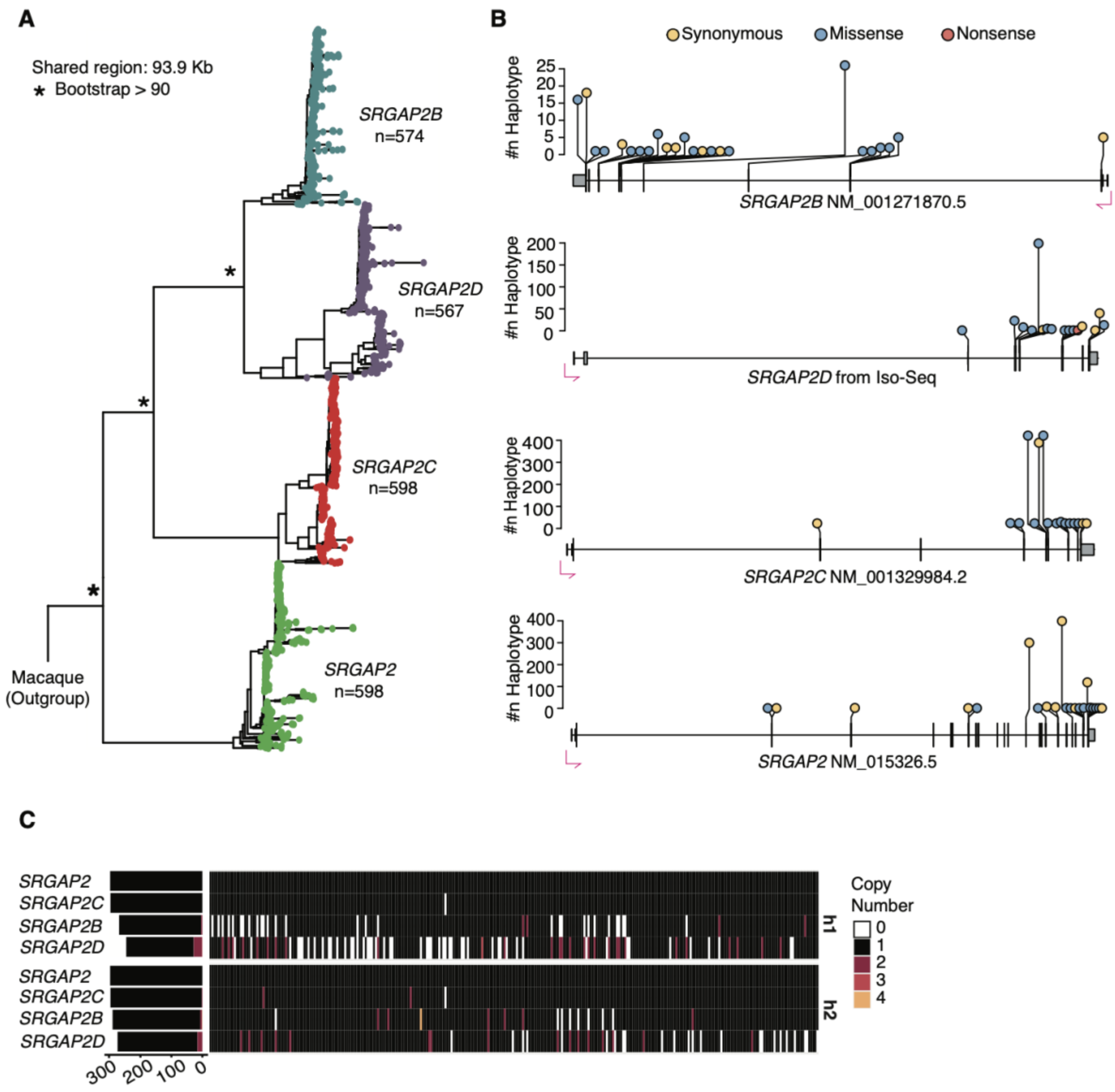


**Supplementary Fig. 14 | Paralog grouping, copy number, and variants for *SRGAP2* gene family.** Same legend as Supplementary Fig. 5.

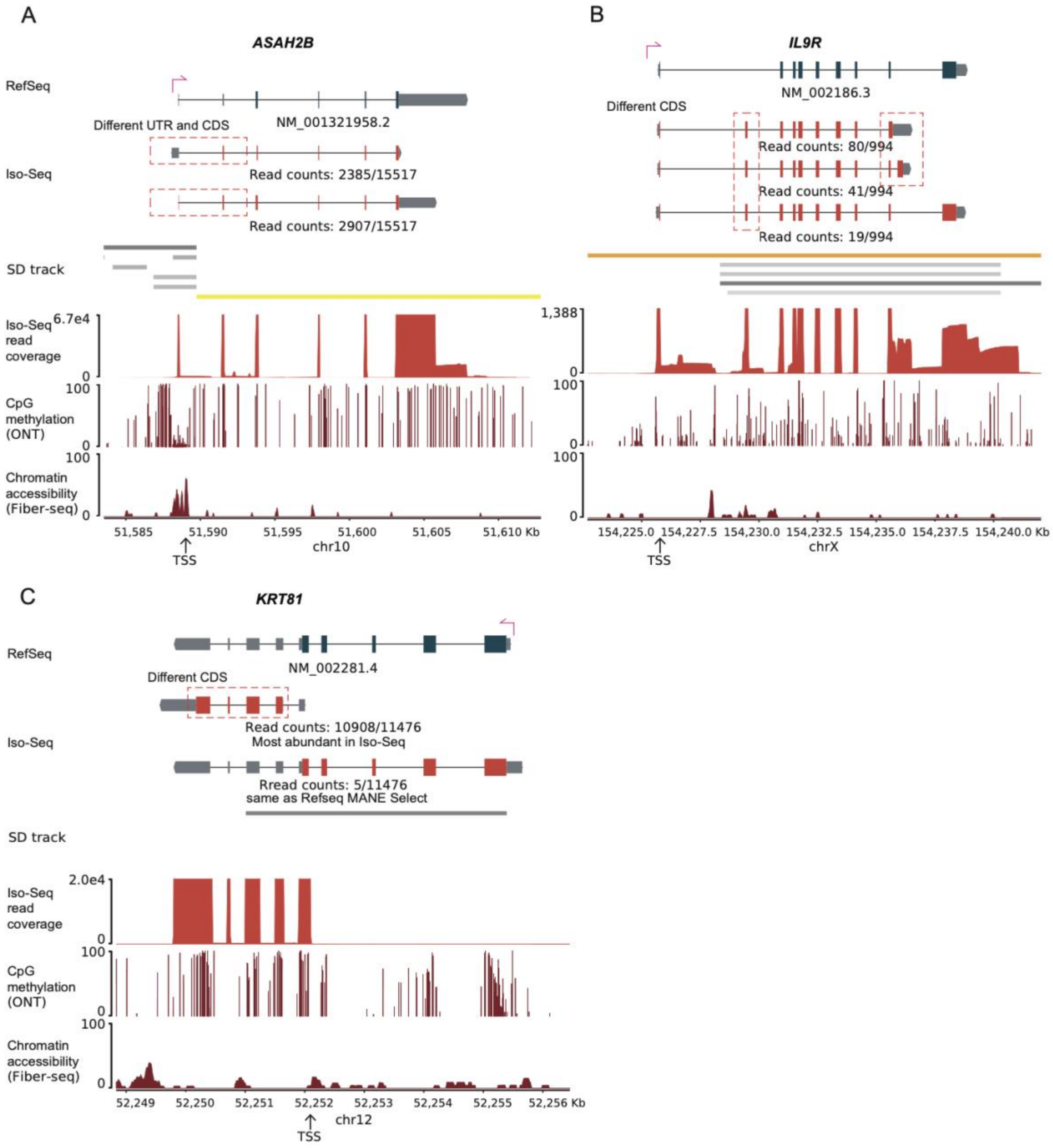


**Supplementary Fig. 15 | Gene models identified using Iso-Seq.** Four genes showed changes in constraint status after their gene models were revised based on Iso-Seq data. Three are shown here: **(A)** *ASAH2B***, (B)** *IL9R*, and **(C)** *KRT81*; *NOTCH2NL2* is shown in Supplementary Fig. 12A. The two or three most abundant gene models are displayed for each gene. The gene model selected for variant identification and classification was determined by jointly considering transcript abundance and read coverage.

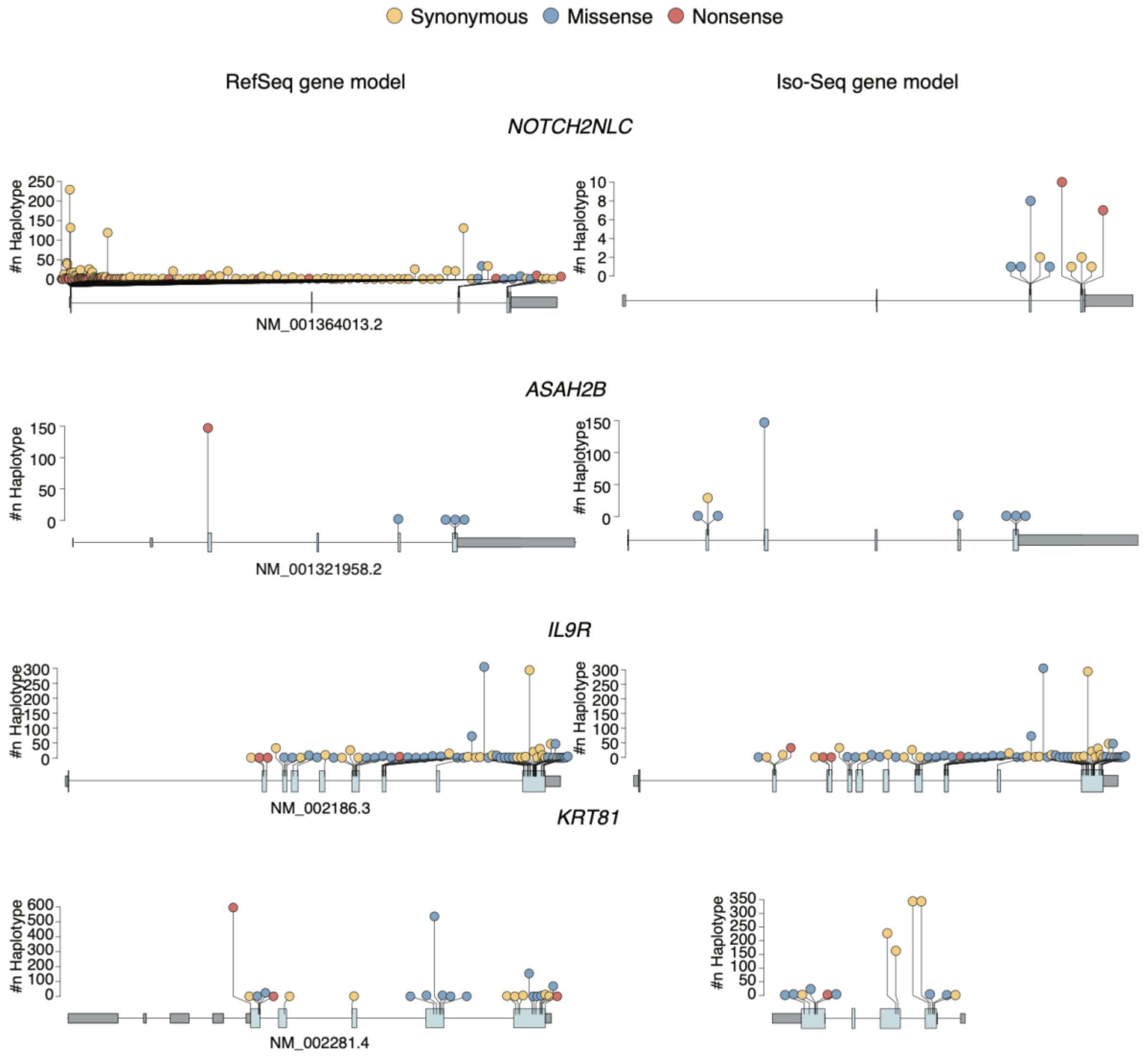


**Supplementary Fig. 16 |** Variants identified in four genes (*NOTHC2NLC*, *ASAH2B*, *IL9R* and *KRT81*) whose constraint status changed after revision of their gene models using Iso-Seq data. Variants are shown based on the RefSeq gene models (left panels) and the revised Iso-Seq gene models (right panels). Amino acid constraint is illustrated by synonymous variants (yellow), amino acid substitutions (blue), and stop-gain variants (red), with variant consequences predicted using VEP.

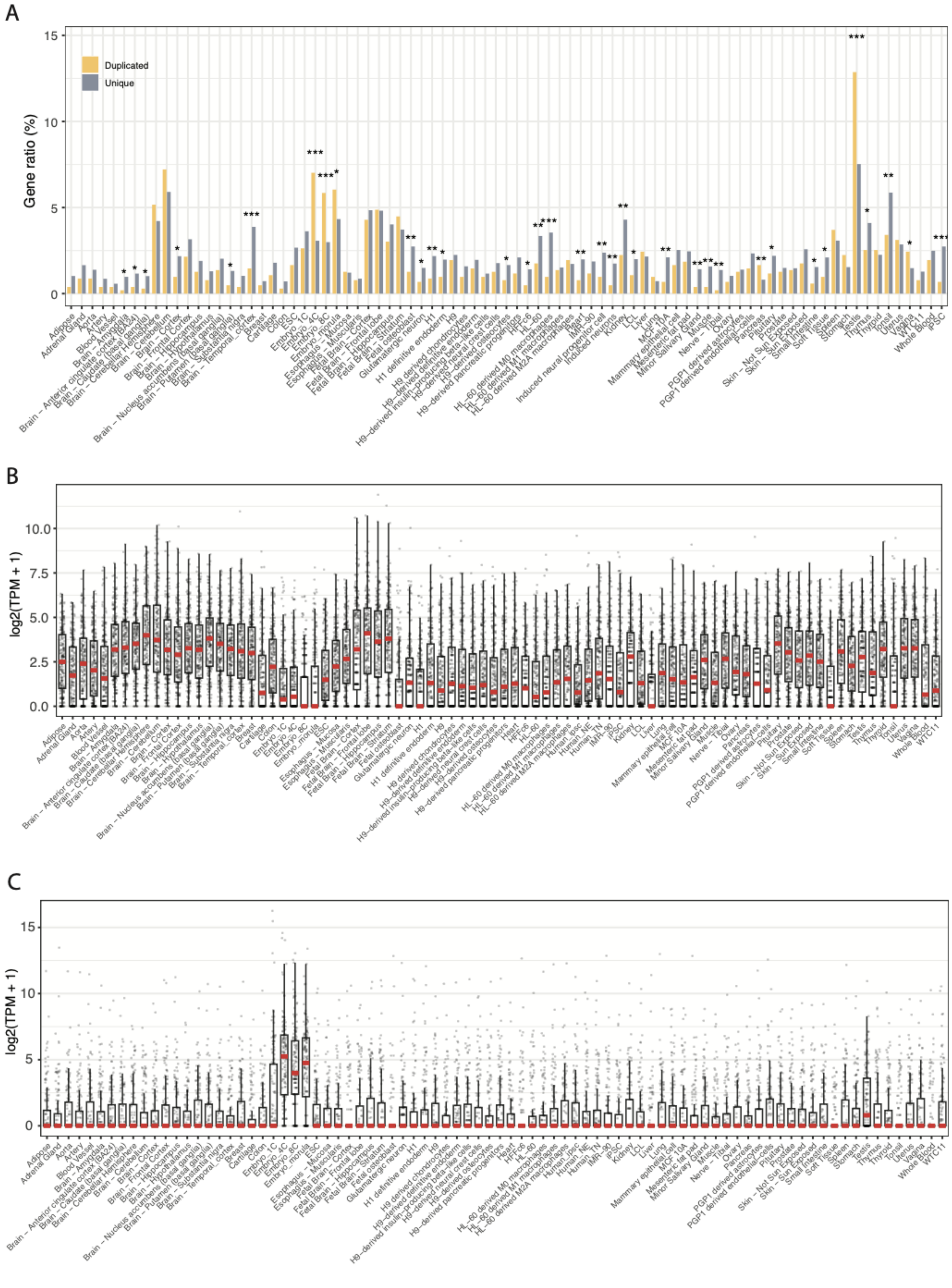


**Supplementary Fig. 17 | Tissue enrichment and expression profiles of duplicated genes. (A)** Tissue-type enrichment comparing SD genes and unique genes. **(B)** Expression levels of 251 duplicated genes are highly expressed in the brain across 83 tissue types. **(C)** Expression levels of 170 duplicated genes highly expressed in embryonic tissues across 83 tissue types.

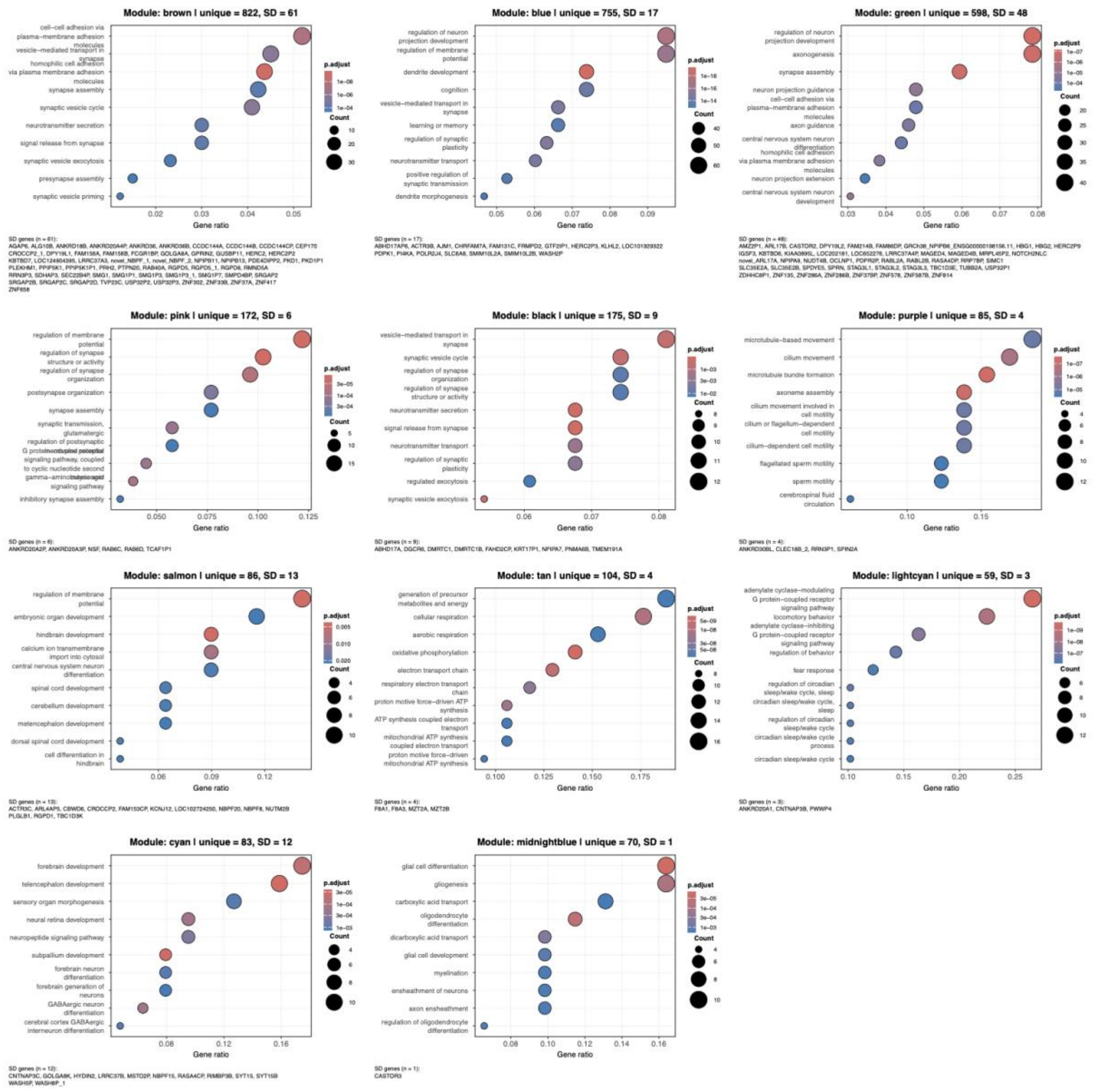


**Supplementary Fig. 18 | GO enrichment analysis of co-expression modules identified by WGCNA for genes highly expressed in the brain.** weighted gene co-expression network analysis (WGCNA) was performed using the expression profiles of 4297 unique genes and 251 SD genes highly expressed in brain tissue, identifying 12 co-expression modules. Genes with a module membership coefficient (kME) greater than 0.5 were retained for downstream analysis, resulting in 3392 unique genes and 202 SD genes across the 12 modules. For each module, unique genes were used as input for GO enrichment analysis to infer the potential biological functions of co-expressed SD genes. Eleven of the twelve modules (containing 178 SD genes) showed significant GO term enrichment (adjusted $p < 0.05$); dotplots display the top enriched biological process terms for each module, with corresponding SD genes listed below each plot.

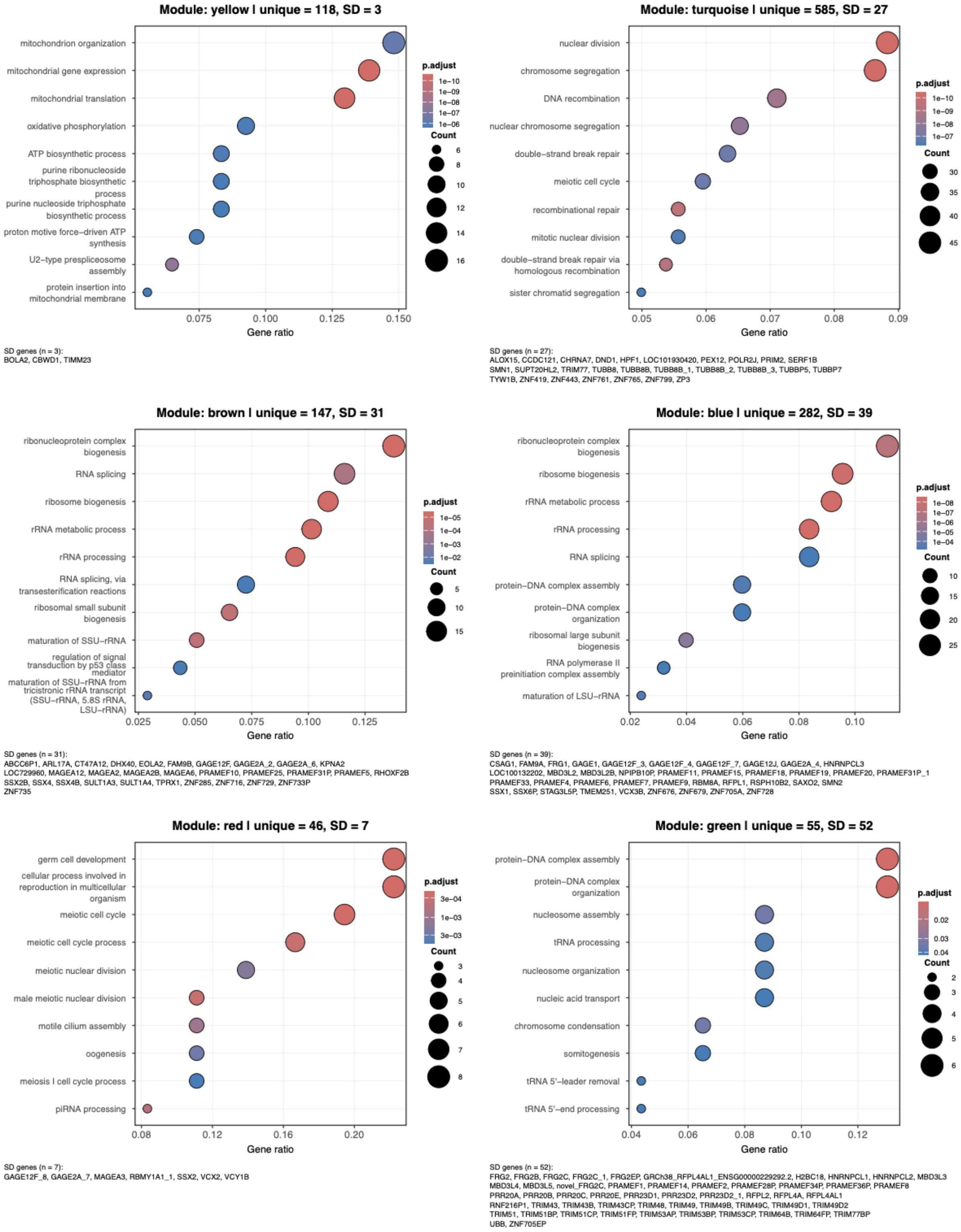


**Supplementary Fig. 19 | GO enrichment analysis of co-expression modules identified by WGCNA for unique and duplicated genes highly expressed in the embryo.** WGCNA was performed using the expression profiles of 1515 unique genes and 170 SD genes highly expressed in embryonic tissue, identifying six co-expression modules. After filtering by kME, 158 SD genes were retained for GO analysis. Same legend as Supplementary Fig. 18.

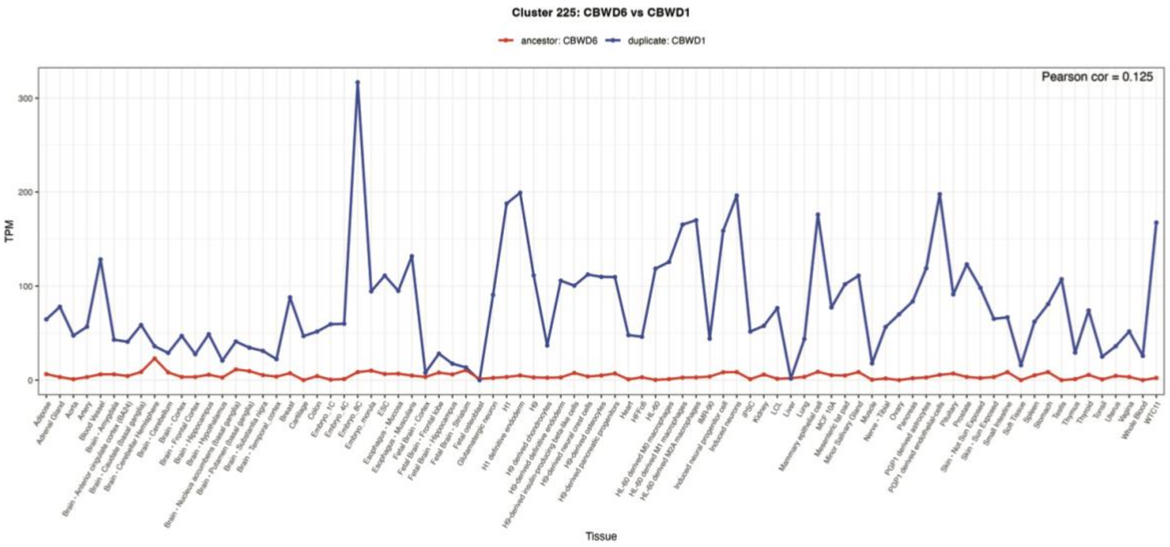


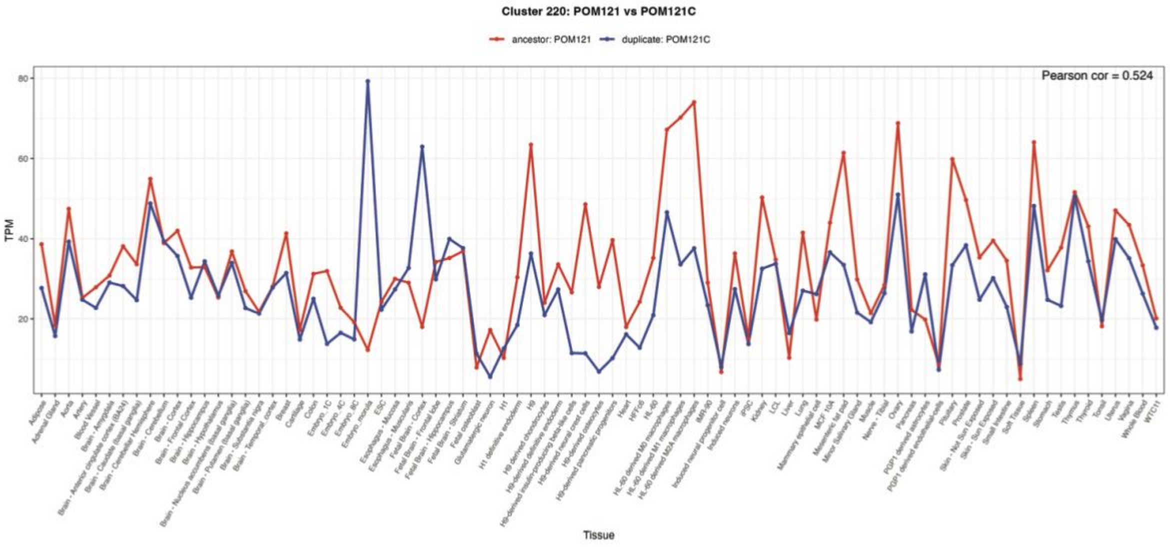


**Supplementary Fig. 20 |** Gene expression of *CBWD6* (ancestor) and *CBWD1* (derived), and *POM121* (ancestor) and *POM121C* (derived) across 83 tissue types.

### Supplementary Tables

**Supplementary Table 1.** 1877 SD genes analyzed in this study, comprising 1811 genes from T2T-CHM13, 24 unannotated genes, and 42 non-CHM13 genes identified from phylogenetic trees.

**Supplementary Table 2.** Paralog-level copy number for each monoclade. For each assembly, the number before the pipe symbol (|) indicates the copy number identified prior to Flagger assembly quality filtering, and the number after the pipe symbol indicates the copy number retained after removing copies within assembly errors. When ancestral and derived genes are grouped within the same monoclade, the ancestral status of that monoclade is assigned as undetermined.

**Supplementary Table 3.** Non-reference SD gene families. Genes that are unique in T2T-CHM13 but duplicated in other human assemblies.

**Supplementary Table 4.** 583 Iso-Seq datasets used for gene model validation and gene expression analysis.

**Supplementary Table 5.** Curated gene annotations supported by long-read Iso-Seq transcript evidence.

**Supplementary Table 6.** Copy number and amino acid variant constraint summary for SD genes, including population-level copy number fixation and LOF variant statistics for each paralog. The column "hap_constraint" indicates constraint based on haploid copy number and amino acid constraint, and "dip_constraint" indicates constraint based on diploid copy number and amino acid constraint.

**Supplementary Table 7.** Highly expressed SD genes in brain, embryo, and testis.

**Supplementary Table 8.** Expression pattern between ancestral and derived SD genes.
